# A shotgun approach for highly multiplexed mammalian metabolic engineering

**DOI:** 10.1101/2025.07.08.663766

**Authors:** Julie Trolle, Sofia Sessa, Tori Rodrick, Drew R Jones, David Fenyö, Sudarshan Pinglay, Jef D Boeke

## Abstract

Mammalian metabolic engineering is critical to advancing basic biology, bioproduction, and cell therapy. However, as pathway complexity increases, so does the size of both the combinatorial design space and the DNA constructs required, rendering unbiased screens intractable. To address this, we developed Shotgun Genetic Engineering (SGE), a scalable approach that exploits the ease of delivering many barcoded small constructs—rather than a single large one—into mammalian cells. This allows each cell to serve as an independent experiment, carrying a unique synthetic metabolic pathway that explores combinations of gene content, stoichiometry, and organellar localization. Functional pathways are identified by sequencing barcodes from cells exhibiting the desired phenotype. Using SGE, we screened millions of pathway combinations to engineer essential amino acid biosynthesis in two mammalian cell lines (CHO and Jurkat), achieving near-wild-type growth in valine-free medium and, for the first time, enabling isoleucine prototrophy in CHO cells. Successful solutions favored mitochondrial localization and required integration of 23–52 kb of synthetic DNA—lengths that are impractical to screen by conventional methods. The resulting datasets are compatible with machine learning frameworks, positioning SGE as a powerful platform for decoding and engineering complex biosynthetic traits in mammalian systems.

## INTRODUCTION

The ability to exert precise and predictable control over cellular behavior is a central goal of synthetic biology. In metabolic engineering, this control is often achieved by coordinating the activity of multiple transcription units (TUs), each consisting of a gene and its regulatory elements. Synthetic metabolic pathways are frequently constructed by transplanting coding sequences from diverse species into orthogonal hosts, thereby endowing them with new metabolic capabilities. This approach has yielded notable successes, including the development of cell-based alternatives to industrial manufacturing processes, the stabilization of pharmaceutical supply chains previously dependent on seasonal plant sources, and the capture of atmospheric carbon dioxide to generate industrially valuable biomass^1–3^. While these strategies have been extensively demonstrated in microbial chassis, examples of metabolic engineering at this pathway scale in mammalian cells remain rare^4^, despite broad recognition that improved control of mammalian cell function could enhance cell therapies, help produce increasingly complex biologics, and the improve the scalable manufacture of viral vectors^5–7^. Modern DNA synthesis technologies now make it possible to incorporate sequences from across the phylogenetic tree or *de novo* designed sequences, vastly expanding the functional landscape accessible to mammalian cells.

Despite this promise, the limited examples of mammalian metabolic engineering can in part be attributed to the lack of foundational knowledge needed to guide metabolic pathway design. Given a target metabolic pathway and a target host cell type, one is faced with a series of complex design decisions - including the selection of coding sequences (CDSs), the configuration of regulatory elements, the intracellular localization of each encoded protein, and the relative expression levels and stoichiometry of each TU required for optimal function (**Figure 1A**). These variables create a vast combinatorial design space, yet there is little mechanistic understanding to guide optimal choices.

**Figure 1.**
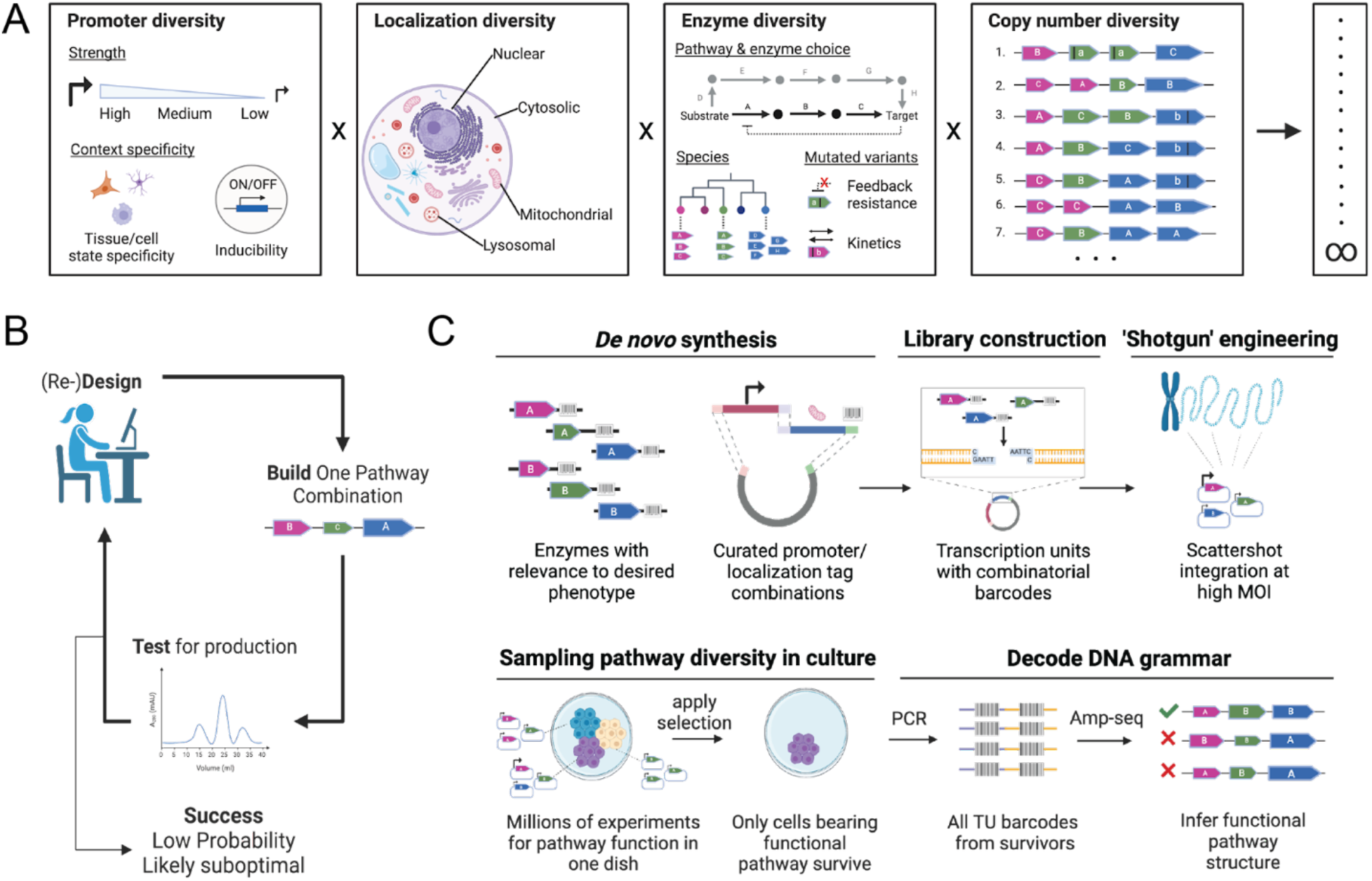
**Shotgun Genetic Engineering expands DNA search space to sample many metabolic pathway solutions in parallel** A. Metabolic engineering benefits from increasing diversity across the different modules, which make up transcription units, including promoters, organellar localization signals, enzymatic function, stoichiometry, and copy number. B. Classical Design-Build-Test Frameworks in which pathways are tested in succession are inefficient in mammalian cells and other host chassis, which exhibit long doubling times or difficulties associated with engineering/delivering long DNAs that encode more complex biosynthetic pathways. C. Shotgun Genetic Engineering increases the scale of metabolic engineering by delivering barcoded TUs at high multiplicity of infection (MOI) such that each cell carries a unique pathway combination - thereby sampling much more pathway diversity. Cells displaying the desired phenotype can be sequence to reveal TU combinations that resulted in function.

In fast-growing, genetically tractable microbial systems such as *Escherichia coli* (20-minute doubling time) or *Saccharomyces cerevisiae* (90-minute doubling time), the lack of mechanistic knowledge can often be mitigated through the design-build-test (DBT) framework, where metabolic pathways are rapidly constructed, tested, and iteratively refined until satisfactory performance is achieved^8,9^ (**Figure 1B**). Applying similarly complex metabolic engineering strategies in mammalian cells is far more challenging, primarily due to much longer doubling times (typically over 20 hours), which drastically slows the pace of iteration.

This limitation becomes increasingly severe as pathway complexity grows or as multiple functions are integrated into a single cell. While the final implementation of such complex functions may rely on recent advances in the synthesis and delivery of large DNA constructs to mammalian cells >100kb^10^, these approaches are poorly suited for high-throughput screening, as delivering large DNA constructs into mammalian cells is highly inefficient. Modular combinatorial strategies, such as CLASSIC, offer a solution by enabling screening ∼10^5^ constructs at the ∼10 kb scale^11^ (2-3 TUs); however, there is a strong inverse relationship between DNA size and delivery success - it becomes exponentially more difficult to deliver larger constructs, rendering brute-force screening of complex synthetic designs impractical^12^.

To overcome the limitations of experimental screening in mammalian systems, some metabolic engineering approaches incorporate *in silico* predictions of metabolic networks to prioritize genetic interventions before testing in the lab^11,13^. Sophisticated transcriptomic databases and metabolic models have been developed for commonly used mammalian cell types, enabling the identification of epistatic metabolic functions and the design of complex multiple-knockout cell lines that improve bioprocessing and manufacturability of antibodies^14–16^. However, these models are typically trained on observational data shaped by natural evolution and are optimized for a narrow range of species and cell types commonly used in biomanufacturing. As a result, they are poorly suited for generative design, including scenarios involving the introduction of heterologous or entirely synthetic TUs. What is needed is *synthetic data*: systematic, large-scale metabolic perturbations that can be used to train machine learning models capable of predictive and design-oriented applications^17^.

Here, we introduce Shotgun Genetic Engineering (SGE), a multiplexed approach to mammalian metabolic engineering that enables rapid discovery of complex pathway solutions and the creation of large-scale synthetic datasets for training predictive models. SGE leverages high-multiplicity integration of barcoded individual TUs from a pooled library, allowing distinct pathway variants to be randomly assembled within each cell. Because small DNA constructs are exponentially easier to synthesize, assemble, and deliver, this design turns each cell into an independent experiment and enables millions of combinations to be screened in parallel. By selecting for cells with desired phenotypes and sequencing their associated barcodes, SGE bypasses the limitations of sequential design-build-test cycles and dramatically accelerates discovery timelines (**Figure 1C**).

As a proof-of-concept, we apply SGE to screen millions of pathway combinations for engineering two essential amino acid biosynthetic pathways in mammalian cells. In prior work, we enabled CHO cells to proliferate without valine by introducing four *E. coli* genes, achieving a valine-free doubling time of 4.3 days^18^. However, we were unable to confer isoleucine independence, despite overlapping biosynthetic steps. Here, we use SGE to both reconstitute and optimize valine prototrophy in CHO cells, yielding clones with a near-wild-type doubling time of just 1.1 days. We also discover, for the first time, a functional six-gene solution enabling isoleucine prototrophy in CHO cells. In both cases, we saw strong signatures favoring mitochondrial localization of biosynthetic enzymes for optimized pathway functionality, indicating that organellar localization is an underexplored variable in mammalian metabolic engineering. Extending this approach, we introduce valine prototrophy to a human T lymphocyte (Jurkat) cell line - an initial demonstration of metabolic pathway expansion in immune cells, with potential applications in engineering stress-resilient cell therapies. Finally, we show that SGE enables training of a machine learning classifier to identify genetic features predictive of pathway function. Together, these results establish SGE as a scalable framework for pathway-scale metabolic engineering in mammalian cells.

## RESULTS

### Design of Shotgun Genetic Engineering

A central premise of SGE is that individual transcriptional units (TUs) are exponentially easier to synthesize, assemble, and deliver than large multi-gene constructs. By introducing a pooled library of individually barcoded TUs at high multiplicity of infection (MOI), each cell randomly assembles a unique combination of TUs that, within the cellular environment, function together as a pathway, effectively turning each cell into a self-contained experiment. This approach eliminates the need for laborious *in vitro* pathway construction or inefficient delivery of large DNA constructs, enabling scalable exploration of vast pathway space in a single multiplexed experiment.

SGE further exploits the modularity of TUs to maximize combinatorial diversity. A curated set of genes hypothesized to influence a phenotype of interest is selected, and each coding sequence (CDS) is assigned a unique set of barcodes. These CDSs are synthesized with flanking Type IIS restriction sites to enable directional insertion into a pooled set of pre-assembled vectors carrying a diversity of barcoded promoters. To investigate the role of subcellular localization, promoters can be paired with N-terminal organellar localization signals (OLSs), which receive additional identifying barcodes. These components - CDS, promoter, and optionally OLS - are assembled in a single-pot Golden Gate reaction into lentiviral-compatible expression vectors.

Critically, each of these elements is fully modular and exchangeable: the library can be expanded to include arbitrarily large sets of CDSs, promoter-OLS pairs, host factor perturbations, or alternative delivery backbones (e.g., transposons or integrases). Each assembly vector places the promoter and CDS barcodes within ∼100 bp of each other in the 3′ UTR, allowing efficient co-recovery via a single PCR amplicon.

The pooled library is packaged into lentivirus and delivered to cells at high MOI. Lentiviral integration adds an additional layer of diversity through variable TU copy number and stoichiometry, yielding a heterogeneous population in which each cell encodes a unique configuration of the pathway. Cells are then subjected to a selection pressure that favors the outgrowth of functional pathway configurations. Clones exhibiting a selective advantage can be phenotyped, and the underlying TU composition inferred by sequencing the linked barcodes (**Figure 1C**). Although in this study we focus on growth-based selection, the selective pressure in SGE can be applied through a variety of mechanisms - including biosensors, fluorescence-based sorting, or other functional readouts - depending on the phenotype of interest. This architecture makes SGE uniquely suited for scalable, pathway-level exploration of genotype–phenotype relationships in mammalian cells.

### SGE Valine/Isoleucine Library Curation and Construction

To confirm the feasibility of SGE as a metabolic engineering approach, we focused on reconstituting essential amino acid (EAA) biosynthesis in mammalian cells. This challenge offers a stringent test case: EAA biosynthesis was lost in the mammalian lineage over 500 million years ago, and mammals must obtain nine amino acids from the environment. In contrast, many microorganisms retain the capacity to synthesize these compounds^19,20^. Engineering EAA pathways into mammalian cells therefore presents a powerful model system to explore the boundaries of metabolic rewiring given its complexity, biotechnological value and experimental tractability. Critically, this system offers a simple and effective selection strategy: cells are grown in the absence of the target amino acid, and only those with functional biosynthetic pathways are able to grow^18^.

We initially set out to reconstitute a valine biosynthetic capacity (prototrophy) in Chinese Hamster Ovary (CHO) cells using SGE, building on prior work in which we had engineered CHO cells using rational design principles^18^. In the earlier study, we found that *E. coli*-derived *ilvN, ilvB, ilvC,* and *ilvD*, were sufficient to confer valine biosynthetic capacity to CHO cells when encoded at a 1:1:1:1 stoichiometry, using 2A ribosomal-skipping peptide sequences (**Figure 2A**). However, this same gene set was insufficient to confer isoleucine biosynthesis to CHO cells despite its overlap in enzymatic steps with valine biosynthesis (**Figure 2B)**. We hypothesized that this might be due to (i) insufficient availability of an isoleucine pathway-specific substrate, 2-oxobutanoate, or (ii) suboptimal catalytic and feedback inhibition properties of the acetohydroxy acid synthase I (AHAS I) complex encoded by catalytic subunit, *ilvB*, and regulatory subunit, *ilvN*.

**Figure 2.**
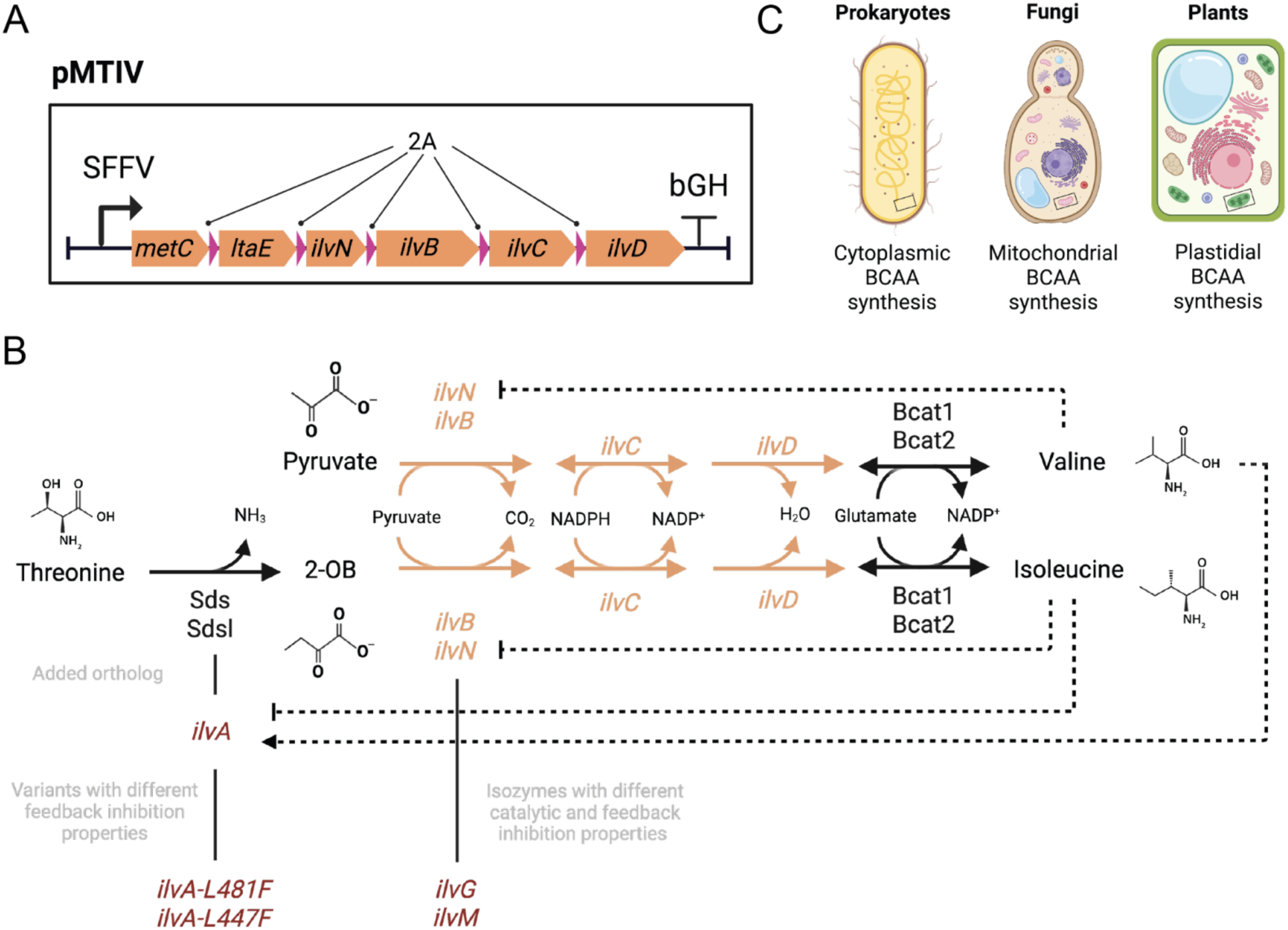
**Engineering branched-chain amino acid biosynthesis in mammalian cells.** A. pMTIV pathway designed in previous study^18^ to successfully confer valine prototrophy to CHO cells. All genes are derived from *E. coli* and encoded in a single open reading frame using 2A ribosomal-skipping peptide sequences. *ilvN, ilvB, ilvC,* and *ilvD* are the subset of genes relevant to valine biosynthesis and were shown to be sufficient to enable valine prototrophy in CHO. B. Pathway map detailing enzymatic steps missing to enable valine/isoleucine biosynthesis in mammalian cells. Black arrows constitute native mammalian genes. Orange arrows constitute *E. coli* enzymes previously imported into CHO to enable valine biosynthesis ^18^. Gene names listed in red are other *E. coli*-derived genes hypothesized to be helpful towards conferring biosynthetic function and which were included in the SGE library. Valine biosynthetic pathway intermediates: pyruvate, (S)-2-acetolactate, (R)-2,3-dihydroxy-3-methylbutanoate, 2-oxoisovalerate, valine. Isoleucine biosynthetic pathway intermediates: threonine, 2-oxobutanoate, (S)-2-aceto-2-hydroxybutanoate, (R)-2,3-dihydroxy-3-methylpentanoate, (S)-3-methyl-2-oxopentanoate, isoleucine. C. Branched-chain amino acid (BCAA) pathways are localized to different intracellular compartments across the phylogenetic tree.

We therefore constructed a library containing *ilvN, ilvB, ilvC,* and *ilvD* as the basis of our library while adding 5 additional genes with the aim of building upon the valine biosynthetic phenotype to potentially discover a biosynthetic pathway composition that would enable prototrophy for isoleucine (**Figure 2B**; **Table 1**). Specifically, we added *E. coli*-derived *ilvA* in the interest of increasing the availability of 2-oxobutanoate as well as two mutated variants of *ilvA* with partial (*ilvA-L481F*) or complete (*ilvA-L447F*) insensitivity to negative feedback inhibition mediated by isoleucine itself. We further added *ilvG* and *ilvM*, isozymes of *ilvB* and *ilvN*, respectively. Together, *ilvG* and *ilvM* form an acetohydroxy acid synthase complex (AHAS II), similar to that formed by *ilvB* and *ilvN* (AHAS I) but with different catalytic properties - most notably a pronounced (57 to 180-fold) preference for isoleucine biosynthetic substrate, 2-oxobutanoate, over the shared isoleucine/valine biosynthetic substrate, pyruvate^21,22^. AHAS II is furthermore insensitive to feedback inhibition by isoleucine and valine, unlike AHAS I, which responds to inhibition by both amino acids. Last, a GFP CDS was included as a control to allow us to monitor behavior of a passenger TU.

**Table 1.** Coding sequences included in the SGE library for conferral of isoleucine and valine biosynthesis to mammalian cells.

| Pathway Step | (Holo)enzyme Name | Gene | Other* |
| --- | --- | --- | --- |
| 0 | Threonine deaminase | *ilvA* | • Sensitive to negative feedback inhibition by isoleucine 45<br>• Sensitive to positive feedback activation by valine 46 |
|  | Threonine deaminase | *ilvA-L447F* | • Completely insensitive to negative feedback inhibition by isoleucine 47,48 |
|  | Threonine deaminase | *ilvA-L481F* | • Partially insensitive to negative feedback inhibition by isoleucine 48,49 |
| 1 | Acetohydroxy acid synthase I (AHAS I) | *ilvB* | • Catalytic subunit<br>• Active in absence of a regulatory subunit but at lower activity.<br>• R = 1 21<br>• Used in pMTIV pathway |
|  |  | *ilvN* | • Regulatory subunit<br>• Mediates AHAS I sensitivity to negative feedback inhibition from valine, isoleucine, and leucine 21<br>• Used in pMTIV pathway |
|  | Acetohydroxy acid synthase II (AHAS II) | *ilvG* | • Catalytic subunit and isozyme of *ilvB*<br>• Active in absence of a regulatory subunit but at lower activity<br>• R = 57-180 21,22 |
|  |  | *ilvM* | • Regulatory subunit and isozyme of *ilvN*<br>• Not sensitive to feedback inhibition 21<br>• Can non-specifically activate *ilvB* to 100% of AHAS I catalytic activity and pyruvate dependence to form a hybrid holoenzyme without sensitivity to feedback inhibition 26 |
| 2 | Ketol-acid reductoisomerase | *ilvC* | Used in pMTIV pathway |
| 3 | Dihydroxy-acid dehydratase | *ilvD* | Used in pMTIV pathway |
| N/A | N/A | *GFP* | Included in library as 'passenger' TU control |

\*Aside from *GFP*, all genes were from *E. coli*. R denotes an isoleucine / valine specificity ratio as outlined in 21

We included two constitutive promoters in the library, EF1a (strong) and PGK (medium). Each promoter appeared either standalone or paired with a mitochondrial targeting sequence (MTS), and each version was assigned 3 unique barcodes. Genes of interest were codon optimized, synthesized, and each gene was also assigned 3 unique barcodes for a total of 9 barcodes per TU to allow added resolution in differentiating unique integration events. We assembled the library using golden gate (Methods) and performed amplicon-seq to monitor distribution of transcription units prior to viral packaging. There was no greater than a 2-fold difference in read count between any two TUs within the library indicating that the library distribution was well-balanced with regard to component parts (**Extended Data 1A**). The library was packaged in lentivirus and 2.42M CHO cells were infected at an MOI of 8.8 as estimated by qPCR following infection (**Extended Data 1B)**.

In addition to direct TU modules (promoter, CDS, OLS), each integrated cassette further encompassed a woodchuck hepatitis virus post-transcriptional regulatory element (WPRE), anti-repressor elements (AREs), and lox sites to enable excision of TUs using Cre recombination. Given an average integrated cassette size of 4092 bp, this corresponds to 36 kb of synthetic DNA sampled per infected cell or 8.7 x 10^10^ bp sampled across the infected, diversified population (CHO-Div). Of this, TU modules averaged 2146 bp, corresponding to 19 kb of TU modules sampled per infected cell or 4.6 x 10^10^ bp sampled across the CHO-Div population. Post-infection, we performed amplicon-seq to monitor TU distribution and observed underrepresentation of EF1a promoters both with and without MTS (**Extended Data 1A**), consistent with reports that use of multiple strong promoters in lentiviral vectors can cause transcriptional interference^23–25^. We also found lesser representation of *ilvA-L447F* (2.6% of reads) and *ilvG* (1.2% of reads) compared to other CDSs.

### Monte Carlo simulation to estimate scale of diversity introduced

This prompted us to perform a Monte Carlo simulation in order to determine whether TU combinations involving these CDSs were likely to be captured in our infection experiment. We simulated randomized choices of TUs based on the barcode distribution measured for CHO-Div at a range of MOIs between 2 and 14 assuming a Poisson distribution for integration events. At an MOI of 8, the simulation indicated that we would observe 1.56M different pathway combinations across the 2.42M infected cells with 12.4% of cells encompassing the gene content of our valine biosynthetic pathway (*ilvN, ilvB, ilvC, ilvD*) while 2.0% of cells would have all 4 genes localized to the cytoplasm and 1.6% would have all 4 genes localized to the mitochondria (**Extended Data 1C**). We reasoned this likely constituted sufficient presence of true pathway solutions for us to discover functional valine biosynthetic pathways post-selection and proceeded to selection on a valine-restricted medium.

### Reconstitution and Optimization of Valine Prototrophy in CHO

CHO-Div was subject to reduced (4.25 nM) valine medium to favor outgrowth of cells carrying combinations of TUs that together comprised biosynthetic pathways for valine (**Figure 3A**). In parallel, we subjected a control population (CHO-GFP) that had been transduced with a control virus carrying only *GFP* to the same selection regimen. After 15 days in low valine medium, we found clonal outgrowths of cells in the experimental population (CHO-Val) but not in the control CHO-GFP population (**Figure 3B)**. We picked 16 CHO-Val clones and passaged each individually in a valine-replete (170 nM) medium prior to characterization via an amino acid dropout assay and amplicon-seq to determine the combination of TUs present. As a proxy for cell number, metabolic activity was measured using a resazurin-based reagent, PrestoBlue, and valine biosynthetic function was estimated by measuring the ratio of valine-free to valine-replete metabolic activity (Val Score). We considered a clone valine prototrophic if its Val Score was at least 2X that of the parental control. 100% of selected clones (16/16) met this threshold (**Figure 3C**). To verify that this functional assay accurately captures valine prototrophic function, we further characterized two clones exhibiting a Val Score > 0.75, CHO-Val-D1 and CHO-Val-D3. We cultured both clones in valine-free medium over 10 days, measuring their absolute growth rate. While control cells died off by day 7 in valine-free medium, clones CHO-Val-D1 and CHO-Val-D3 instead proliferated over the 10 days of valine-free culture at rates of 1.14 and 1.05 days/doubling, respectively, (**Figure 3D**) corresponding to 83% and 80% of their growth rates in valine-replete medium **(Extended Data 2A**). This is a significant improvement in valine-free growth rate compared to the 3.77 days/doubling growth rate exhibited by pMTIV cells that we engineered using rational design in our previous study^18^, which corresponded to just 26% of their valine-replete growth rate.

**Figure 3.**
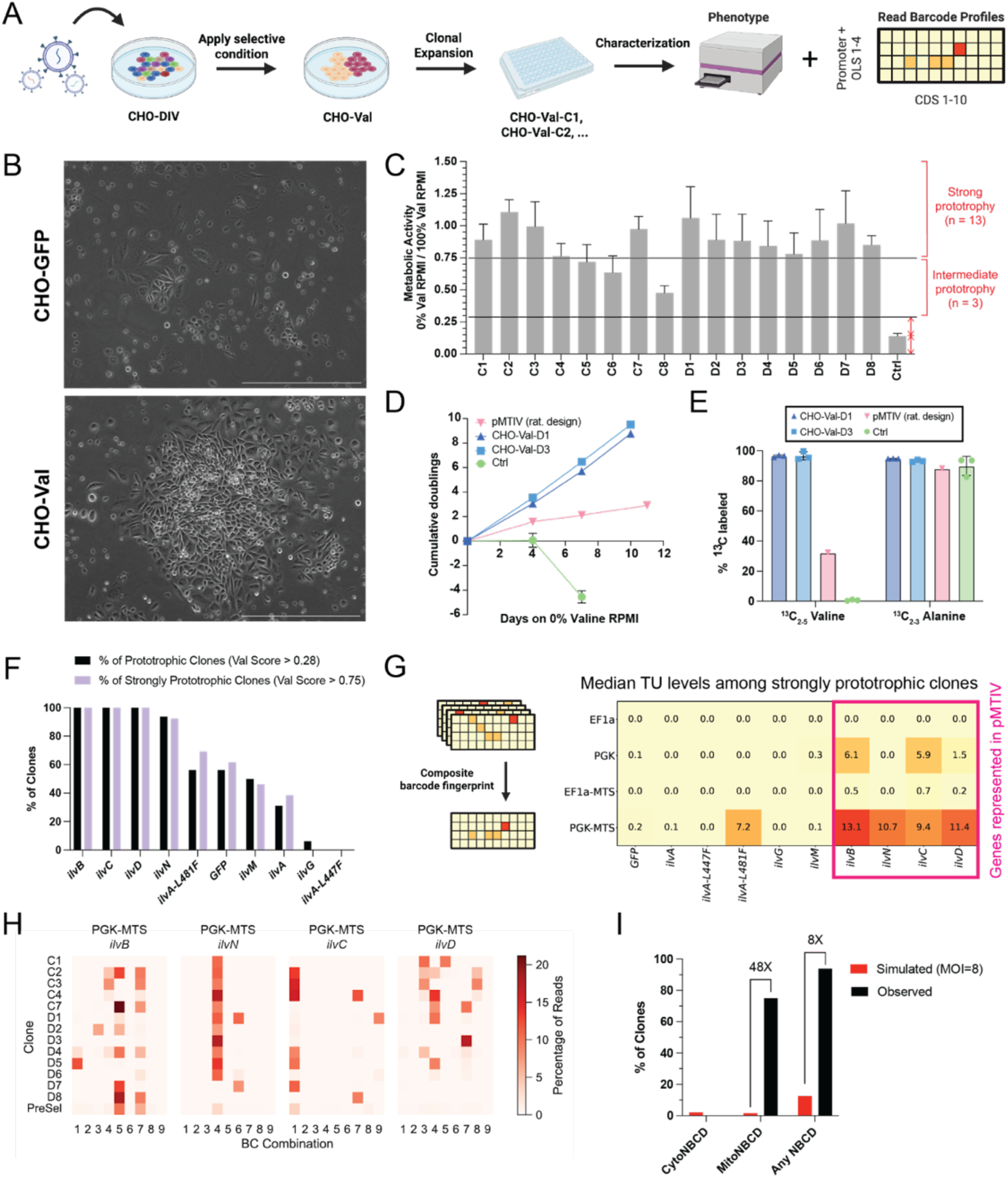
**Reconstitution of valine biosynthesis in CHO to validate and benchmark Shotgun Genetic Engineering against a rational design approach** A. Schematic detailing SGE workflow from infection to phenotyping and barcode readout for adherent CHO cells. B. Representative images of clonal outgrowths after 15 days of selection on low valine RPMI medium from populations infected with either GFP or SGE library. Scale bar corresponds to 500µm. C. PrestoBlue amino acid dropout assay of 16 individual clones selected on low valine RPMI and the parental control cell line (Ctrl). Each clone was measured in triplicate wells after growth in valine-free medium for 3 days and compared to triplicate wells cultured on valine-replete medium for 3 days to generate a ‘Val Score’. Valine prototrophy was defined as a 2X improvement in Val Score relative to a non-prototrophic parental cell line. Strong valine prototrophy was arbitrarily defined as a Val Score of 0.75. Error bars represent standard deviation across triplicate wells. D. Growth curve of Clones CHO-Val-D1 and CHO-Val-D3 cultured in valine-free RPMI and compared to the parental cell line as well as a pMTIV cell line engineered in a previous study using rational design principles. Error bars represent standard deviation across triplicate wells. E. Endogenous ^13^C valine labeling in cells cultured on RPMI with ^13^C_6_ glucose and ^13^C_3_ sodium pyruvate. Error bars represent standard deviation across triplicate wells for Clones CHO-Val-D1, CHO-Val-D3, and parental control cells. pMTIV is a single replicate from previous study^18^. F. Monitoring CDS presence across all valine prototrophic clones (Val Score > 0.28) and across all strongly valine prototrophic clones (Val Score > 0.75) G. Composite barcode ‘fingerprint’ comprised by median normalized read count across all strongly valine prototrophic clones (Val Score > 0.75) H. Heatmap depicting abundance of TUs involved in conferring valine prototrophy and their 9 underlying barcode combinations (3 per promoter/OLS, 3 per CDS) across all strongly valine prototrophic clones (Val Score > 0.75) I. Comparing representation of *ilvN, ilvB, ilvC*, and *ilvD* in clones selected on low valine medium relative to their representation if selected by random chance as determined by a simulated infection at MOI=8.

To further confirm that cells were endogenously synthesizing valine, we cultured CHO-Val-D1 and CHO-Val-D3 in a valine-free ‘heavy’ RPMI medium in which ^12^C glucose was substituted for its fully labeled ^13^C counterpart and to which 2 mM ^13^C sodium pyruvate was added (**Extended Data 2B**). Both clones were cultured in this valine-free heavy medium over 3 passages while control cells were cultured in an equivalent heavy medium but with valine added back to facilitate their growth on ^13^C-labeled valine pathway substrates over the same timeframe. Using LCMS, we confirmed detection of ^13^C_5_-labeled valine in our engineered clones via extracted ion chromatogram, MS1, and MS2 analysis (shown for CHO-Val-D1 in **Extended Data 2C-E**). In quantifying ^13^C-labeling levels of valine across cell lines, we considered only isotopologues of valine with at least 2 carbon-13 atom incorporations as being endogenously biosynthesized given that carbon-13 constitutes approximately 1% of all carbon. Therefore, the natural abundance of ^13^C_1_ valine is 5% (5 carbons x 1% representation) while ^13^C_2_ valine is only expected to represent 0.25% of the total pool (5% x 5%). Both clones exhibited considerably increased ^13^C_2-5_ valine labeling at 96.5% and 96.6% respectively, compared to pMTIV cells at 32.2%, indicating strong improvement of the valine biosynthetic phenotype compared to the rationally designed metabolic pathway (**Figure 3E**). Indeed, ^13^C_2-5_ Valine levels detected in Clones Val-D1 and Val-D3 matched ^13^C_2-3_ Alanine levels, suggesting that the engineered biosynthesis of valine in these clones is on par with the native mammalian cellular ability to biosynthesize the non-essential amino acid alanine, which is also biosynthesized from pyruvate.

To understand the genotypic differences between clones, we performed amplicon-seq to gain insight into the relative distribution of CDSs within all clones including in CHO-Val-D1 and CHO-Val-D3, which appeared to carry 9 and 6 TU integrations, respectively, corresponding to 37 and 23 kb of integrated DNA, respectively, of which 19 and 11 kb made up by TU modules. In both clones, we found barcodes corresponding to *ilvN*, *ilvB*, *ilvC*, and *ilvD*, consistent with the CDS content of the rationally designed pMTIV pathway with CHO-Val-D3 additionally exhibiting barcode signatures for *ilvM*, an isozyme of *ilvN* (**Extended Data 3A**). In order to ascertain which CDSs were essential to conferring valine biosynthetic function to cells, we identified CDSs that were universally present across all prototrophic clones (intermediate + strong; Val Score > 0.28) (**Extended Data 3B**). We found *ilvB*, *ilvC*, and *ilvD* presence in every clone exhibiting prototrophy indicating their necessity in conferring valine biosynthetic function to CHO cells (**Figure 3F**). On the other hand, *ilvN* appears in 94% (15/16) of clones after selection, suggesting that it supports valine biosynthetic function without being strictly necessary. This is consistent with reports that *ilvN* is a regulatory subunit that boosts *ilvB* catalytic rates and substrate specificities but that *ilvB* can perform its catalytic function in the absence of *ilvN*. However, in the one clone which lacked *ilvN*, CHO-Val-D8, cells instead carried its isozyme, *ilvM* (**Extended Data 3C**), suggesting a possible role for a hybrid AHAS holoenzyme *ilvBM*, which has previously been demonstrated to be functional and demonstrates differential catalytic and feedback properties compared to its canonical AHAS I and AHAS II counterparts^26,27^.

We next wanted to determine the optimal promoter and localization context for each CDS and thus looked at the distribution of barcodes corresponding to each CDS across the 4 promoter/OLS combinations, focusing specifically on clones exhibiting strong prototrophy (Val Score > 0.75). Among the 13 strongly prototrophic clones we generated a composite barcode ‘fingerprint’ consisting of the median representation of each of the 40 unique TUs within each sample. Within the composite fingerprint, we examined the universally present CDSs *ilvB*, *ilvC*, and *ilvD*, as well as the near-universally present *ilvN*, and found that each was most strongly favored when expressed from a mitochondrially localized PGK promoter. Together, these elements formed a pathway “solution” that had not been previously constructed using rational design (**Figure 3G**).

Given that each TU has 9 underlying combinatorial barcodes (3 barcodes per promoter/OLS combination, 3 barcodes per CDS), we were able to examine the underlying barcode patterns for each TU within our pathway solution. We found a diversity of barcode patterns underlying each TU among strongly prototrophic valine clones, suggesting that the different clones favored during low-valine selection were derived from multiple independent integration events (**Figure 3H**). Last, we examined how often *ilvN*, *ilvB*, *ilvC*, and *ilvD* were collectively observed to be cytoplasmically or mitochondrially localized in the expanded clones as well as how often they were observed irrespective of intracellular location and compared each to expectation based on simulated data in which TUs were selected at random assuming an MOI=8. We found no observed occurrences of collectively cytoplasmically localized *ilvN*, *ilvB*, *ilvC*, and *ilvD*, while mitochondrial localization of *ilvN*, *ilvB*, *ilvC*, and *ilvD* was dramatically selected for at 48X higher representation than expected by chance (**Figure 3I**). Together, these results demonstrate the utility of SGE for identifying solutions to complex metabolic engineering problems.

### Discovery of metabolic pathway solutions to encode isoleucine prototrophy in CHO

In the interest of discovering combinations of TUs that together comprise a functional isoleucine prototrophic phenotype, CHO-Div cells were further subject to a series of reduced (1.5 nM; 3 nM; 6 nM) isoleucine RPMI conditions. In parallel, we subjected a control CHO-GFP population to the same selection regimen. After 15 days in low isoleucine medium, we found large clonal outgrowths of cells in the experimental populations (CHO-Ile) across all conditions but not in the control CHO-GFP population. Across the 3 isoleucine conditions, we picked 46 clones and passaged each individually in an isoleucine-replete (60 nM) medium prior to performing functional characterization and amplicon-seq. As previously, we estimated the isoleucine biosynthetic capacity of each clone by measuring the metabolic activity of each clone cultured in isoleucine-free medium and relativizing to metabolic activity in isoleucine-replete medium (Ile Score). 41% of clones (19/46) exhibited an improved Ile Score relative to the control (**Figure 4A**). To verify that this functional assay accurately captures isoleucine prototrophic function, we further characterized two clones, CHO-Ile-10-H1 and CHO-Ile-10-H2, which exhibited strong and intermediate prototrophy, respectively. We cultured both clones in isoleucine-free medium over 10 days and measured absolute cell numbers during this time. While control cells died off by day 6 in isoleucine-free medium, clone CHO-Ile-10-H1 proliferated over 10 days in isoleucine-free medium at a rate of 1.69 days/doubling, corresponding to 48% of its growth in isoleucine-replete medium, while CHO-Ile-10-H2 proliferated at a rate of 3.04 days/doubling over the same time frame, corresponding to 33% of its growth rate in isoleucine-replete medium (**Figure 4B; Extended Data 4A**).

**Figure 4.**
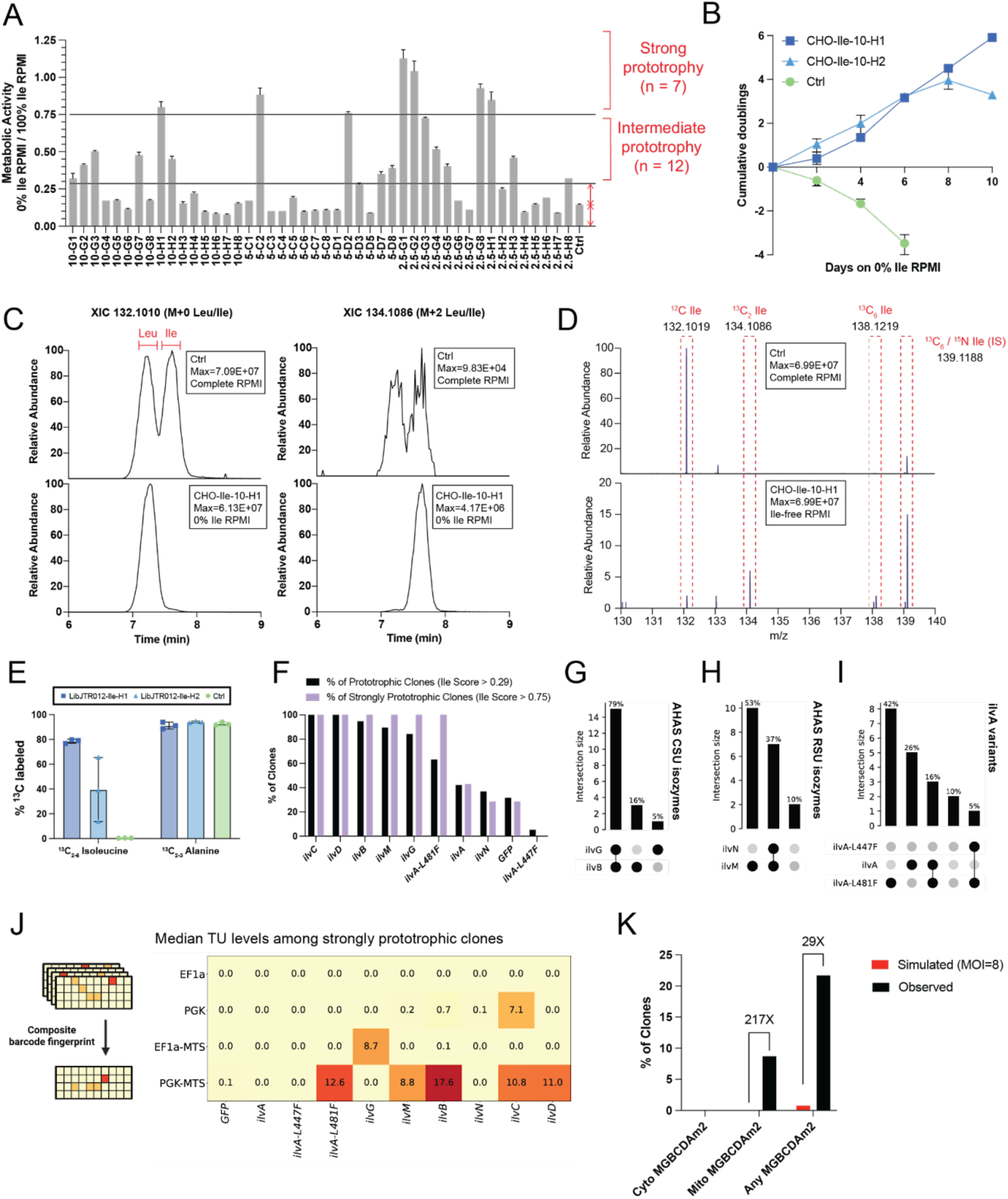
**Discovery of a functional isoleucine biosynthesis pathway in CHO** A. PrestoBlue amino acid dropout assay of 46 individual clones selected on low isoleucine RPMI. Triplicates cultured in isoleucine-free medium for 3 days were compared to triplicates cultured in isoleucine-replete medium for 3 days to generate a ‘Ile Score’. Isoleucine prototrophy was defined as a 2X improvement in Ile Score relative to a non-prototrophic parental cell line. Strong isoleucine prototrophy was arbitrarily defined as Ile Score > 0.75. Error bars represent standard deviation across triplicate wells. B. Growth curve of Clones CHO-Ile-10-H1 and CHO-Ile-10-H2 cultured in isoleucine-free RPMI compared to a non-prototrophic parental cell line. Error bars represent standard deviation across triplicate wells. C. Extracted ion chromatogram showing detection of M+0 leucine and M+2 isoleucine in Clone CHO-Ile-10-H1 cultured on isoleucine-free medium with ^13^C substrates. By comparison, non-prototrophic control cells cultured on isoleucine-replete medium with ^13^C substrates show only M+0 leucine and M+0 isoleucine. D. MS1 peaks corresponding to M+2 and M+6 isoleucine are detected in Clone CHO-Ile-10-H1 E. Quantifying ^13^C isoleucine labeling in cells cultured on RPMI with ^13^C_6_ glucose and ^13^C_3_ sodium pyruvate. Error bars represent standard deviation across triplicate wells. F. Monitoring CDS presence across all isoleucine prototrophic clones (Ile Score > 0.29) and across all strongly isoleucine prototrophic clones (Ile Score > 0.75). G. UpsetPlot showing distribution of AHAS catalytic subunits across 19 isoleucine prototrophic clones. H. UpsetPlot showing distribution of AHAS regulatory subunits across 19 isoleucine prototrophic clones. I. UpsetPlot showing distribution of *ilvA* variants across 19 isoleucine prototrophic clones J. Composite barcode ‘fingerprint’ comprised by median normalized read count across all strongly valine prototrophic clones (Ile Score > 0.75). K. Comparing representation of *ilvN, ilvB, ilvC,* and *ilvD* in clones selected on low isoleucine medium relative to their representation if selected by random chance as determined by a simulated infection that assumed MOI=8.

To further confirm that cells were endogenously synthesizing isoleucine, we cultured CHO-Ile-10-H1 and CHO-Ile-10-H2 in heavy RPMI medium as defined previously. Prototrophic clones of interest were cultured in isoleucine-free heavy medium while control cells were cultured in the same medium but with isoleucine added back in to allow their growth on ^13^C pathway substrates. Using high resolution LCMS, we found accurate mass peaks corresponding to the retention time of both leucine and isoleucine, which share identical molecular weights, at 132.1010 m/z (M+0) in control cells, while only a peak corresponding to leucine was found in engineered clones. At 134.1086 m/z (M+2), the expected labeling signature for endogenous biosynthesis of isoleucine from ^13^C_3_ pyruvate + ^13^C_0_ 2-oxobutanoate, mere background signals of leucine and isoleucine could be seen in control cells while a prominent peak corresponding to isoleucine could be seen in the engineered clones (**Figure 4C**). With MS1 analysis, we observed only M+0 isoleucine and the spiked-in internal standard ^13^C_6_/^15^N_1_ isoleucine in control cells while signatures of both M+0, M+2, and M+6 isoleucine and internal standard ^13^C_6_/^15^N_1_ isoleucine could be found in the prototrophic clones (**Figure 4D**). Using narrow isolation MS2 (+/-0.2 da), we were further able to confirm that the fragmentation signatures for each M+0, M+2, and M+6 as well as internal standard ^13^C_6_/^15^N_1_ isoleucine detected within CHO-Ile-10-H1 were consistent with expectations (**Extended Data 4C**). For clones CHO-Ile-10-H1 and CHO-Ile-10-H2, 78.5% and 39.5% of isoleucine was ^13^C_2-6_, respectively, with the latter exhibiting significant variability in labeling across replicates consistent with its less effective prototrophic function (**Figure 4E**). Together, these clones represent the first demonstration of isoleucine prototrophy in mammalian cells - and, more broadly, the first reconstitution of this metabolic function in the mammalian lineage in over 500 million years.

To understand which TUs were integrated in each clone to confer the isoleucine prototrophic function, we performed amplicon-seq and found significant presence of *ilvB, ilvC, ilvD, ilvG* and *ilvM* in both clones with CHO-Ile-10-H1 additionally carrying *ilvA* and *ilvA-L481F* and CHO-Ile-10-H2 additionally carrying *ilvN* (**Extended Data 5A**). Clone CHO-Ile-10-H1 appeared to carry 13 TU integrations or 52 kb of integrated DNA, while CHO-Ile-10-H2 appeared to carry 10 TU integrations or 41 kb of integrated DNA, of which 26 and 21 kb were directly made up by TU modules, respectively. In order to ascertain which CDSs were essential to conferring isoleucine biosynthetic function to cells, we looked at CDSs that were universally present across all isoleucine prototrophic clones (strong + intermediate) (**Figure 4F**) by again defining a Z-score threshold for CDS presence (**Extended Data 5B**). We found *ilvC* and *ilvD* presence across every prototrophic clone, consistent with expectation as no other isozymes of these enzymes were included in our library. *ilvB* and *ilvM* were present in 95% and 90% of all prototrophic clones, respectively, suggesting their importance in boosting isoleucine biosynthetic pathway functionality without being absolutely necessary to confer prototrophic function (**Figure 4F; Extended Data 5C**). In the one isoleucine prototrophic clone from which *ilvB* was absent, its isozyme *ilvG* was present instead, indicating the necessity of an AHAS catalytic subunit for conferral of isoleucine biosynthetic ability and suggesting that either subunit is sufficient towards conferring isoleucine biosynthetic function (**Figure 4G**). On the contrary, in the two isoleucine prototrophic clones in which *ilvM* did not meet the presence threshold, its isozyme *ilvN* was also not present, suggesting that while *ilvM* is the preferred AHAS regulatory subunit, neither *ilvM* nor *ilvN* are necessary for conferring isoleucine biosynthetic function (**Figure 4H**). Last, we had included 3 *E. coli*-derived variants of *ilvA* in the library, the product of which produces 2-oxobutanoate from threonine with varying feedback inhibition sensitivities. We were curious whether any of these would benefit isoleucine biosynthetic function. Partially feedback insensitive *ilvA-L481F* was present in 63% of clones, natively encoded *ilvA* was present in 42% of clones while completely feedback insensitive *ilvA-L447F* was found in 5% of clones. However, two isoleucine prototrophic clones carried integrations of neither, indicating that bacterial *ilvA* variants are not absolutely necessary for conferral of isoleucine biosynthetic functionality to CHO cells (**Figure 4I**).

We next wanted to determine the optimal isoleucine pathway composition and determined the CDSes that were universally present among strongly isoleucine prototrophic clones (Ile Score > 0.75). Strikingly, 100% of strongly prototrophic clones exhibited barcodes corresponding to *ilvB, ilvG, ilvM* and *ilvA-L481F* in addition to *ilvC* and *ilvD*, suggesting that while not absolutely necessary for isoleucine biosynthetic function, *ilvB, ilvG, ilvM* and *ilvA-L481F* do boost pathway performance (**Figure 4F; Extended Data 4D**). To examine the promoter and localization context for each CDS in strongly prototrophic clones, we again generated a composite barcode ‘fingerprint’ consisting of the median TU level amongst strongly prototrophic clones. We found each CDS to be favored in its mitochondrially localized context with *ilvA-L481F, ilvM, ilvB, ilvC* and *ilvD* favored in their PGK promoter format and *ilvG* in its EF1a promoter format (**Figure 4J**). We examined the 9 underlying barcode possibilities for the favored pathway ‘solution’ and found barcode diversity across the 7 strongly prototrophic clones, suggesting that clones arose from independent integration events (**Extended Data 5E**). We further examined how often *ilvC, ilvD, ilvB, ilvG, ilvM* and *ilvA-L481F* were collectively observed to be cytoplasmically or mitochondrially localized in isolated clones as well as how often they were observed irrespective of intracellular location and compared each to simulated expectation. We found that mitochondrial localization of these TUs was dramatically selected for at 217X expectation while cytoplasmic localization of all 6 CDSs was not observed at all across the 46 clones (**Figure 4K**).

Finally, we looked at CDS representation amongst isoleucine auxotrophic clones (**Extended Data 5F**). Strikingly, we found presence of *ilvB, ilvC*, and *ilvD* across 100% of auxotrophic clones indicating that there was selection for partial metabolic pathways. This is consistent with a model in which infected cells are resource-sharing and metabolic intermediates are available in the medium allowing cells encoding only partial biosynthetic pathways a selective advantage. Such a mechanism would favor CDSs that encode later enzymatic steps in the biosynthetic pathway, which is what we observe (**Extended Data 5G**) with *ilvD* and *ilvC* being most overrepresented relative to simulated expectation followed by *ilvM, ilvG*, and *ilvB*.

### Double isoleucine/valine prototrophy in CHO-Val and CHO-Ile clones

We next cultured all CHO-Val clones and CHO-Ile clones in a double isoleucine/valine dropout medium to ascertain whether we had engineered clones with prototrophy for both amino acids without directly selecting for such a phenotype. Indeed, 37.5% (6/16) of CHO-Val clones and 23.9% (11/46) of CHO-Ile clones surpassed the threshold for double isoleucine/valine prototrophy (2X the metabolic activity of the parental control line; Ile/Val Score > 0.28) although none met the strong double prototrophy threshold (Ile/Val Score > 0.75) (**Extended Data 6A**).

When looking at the genotypes underlying each double prototrophic clone, we found *ilvB, ilvC*, and *ilvD* to be universally present with *ilvM* additionally present in 76.5% of clones (**Extended Data 6B**). However, we noticed that CHO-Val clones and CHO-Ile clones exhibited differential underlying CDS usage suggesting that analyzing both populations on aggregate might conflate signal given that each population was not specifically selected to be doubly prototrophic (**Extended Data 6C**). We therefore focused on parsing out differences within clones amongst the different phenotypes present within each population. Among CHO-Val clones, 6 clones were prototrophic for both valine and isoleucine (Val+Ile+) while 10 were prototrophic for valine only (Val+Ile-). Among CHO-Ile clones, 11 clones were Ile+Val+, 7 clones were Ile+Val- and 21 clones were Ile-Val+ (**Extended Data 6D**). When looking at the gene content underlying each phenotype, CHO-Val clones looked to have similar CDS presence irrespective of phenotype. On the other hand, CHO-Ile cells showed notable differences in *ilvG, ilvA*, and *ilvA-L481F* content (**Extended Data 6E**). For Ile-Val+ CHO-Ile clones, <10% exhibited *ilvG* while the same was true of 100% of Ile+Val-clones, and 75% of Ile+Val+ clones, consistent with *ilvG* specifically supporting isoleucine prototrophy, and consistent with the known catalytic functions of *ilvG* whereby it exhibits 57-180X greater preference for isoleucine biosynthetic intermediates over valine biosynthetic intermediates. On the other hand, *ilvA* was present in just 33% of Ile-Val+ and 33% of Ile+Val+ clones, while 57% of Ile+Val-CHO-Ile clones carried *ilvA*, suggesting that *ilvA* is antagonistic to valine and double valine/isoleucine prototrophic function. Similarly, partially feedback resistant *ilvA-L481F* was present in 29% of Ile-Val+ cells, while the same was true for 57% of Ile+Val-cells and 67% of Ile+Val+ clones, suggesting that *ilvA-L481F* is specifically antagonistic to valine-only prototrophy function and that the feedback resistance of *ilvA-L481F* may strengthen double prototrophic function where *ilvA* only strengthens isoleucine prototrophic function.

Altogether, these results illustrate both the diversity of phenotypes accessible through SGE and the insights such diversity can provide for pathway optimization. Importantly, they underscore a key strength of the SGE framework: the optimal genetic solution for a single metabolic trait (e.g., valine or isoleucine prototrophy) is not necessarily the optimal solution for combined traits (e.g., valine *and* isoleucine prototrophy). Traditional DBTL-based engineering approaches often struggle with such multi-objective tradeoffs, but the scale and combinatorial depth of SGE allows for simultaneous selection across multiple traits, enabling discovery of genotypes that balance or even reconcile conflicting demands across pathways.

### Engineering Valine Prototrophic Human T cells

To examine the versatility of SGE across cell types, we wanted to ascertain whether it was possible to engineer Jurkat cells to become prototrophic for valine and isoleucine. Unlike CHO cells, Jurkat cells grow in suspension, allowing for seamless isolation, expansion and characterization of large numbers of individual clones post-selection either via serial dilution or cell sorting. Using the same lentivirus preparation as employed in previous experiments, we infected 3M Jurkat cells (Jurkat-Div) and found a similar barcode distribution amongst Jurkat-Div cells compared to that previously found for CHO-Div cells (**Extended Data 7A**). The MOI of the infection was estimated by qPCR to be 3.4 (**Extended Data 7B**). Given an average integrated cassette size of 4092 bp, this corresponds to 14 kb sampled per infected cell or 4.2 x 10^10^ bp sampled across the Jurkat-Div population. Of this, TU modules constituted an average of 2146 bp, corresponding to 7.3 kb sampled per infected cell or 2.2 x 10^10^ bp sampled across the entire population. Given the decrease in MOI relative to CHO cells, we again performed Monte Carlo simulations at a range of MOIs (2-8) to determine the likelihood of finding known solutions given the measured barcode distribution (**Extended Data 7B**). At MOI=4, our simulation indicated that we should see 5.9 x 10^5^ different pathways within the 3M infected cells of which 2.31% of cells should carry some combination of *ilvN, ilvB, ilvC*, and *ilvD*, while 0.19% of cells should carry all four TUs localized to the mitochondria and 0.23% all TUs localized to the cytoplasm (**Extended Data 7C**). On the other hand, only 0.06% of Jurkat-Div cells were expected to carry any combination of our newly discovered isoleucine biosynthetic pathway consisting of *ilvM, ilvG, ilvB, ilvC, ilvD* and *ilvA-L481F* with 0.00% of cells expected to carry entirely mitochondrially or entirely cytoplasmically localized versions of all 6 genes (**Extended Data 7D**). We reasoned that we were likely to discover valine prototrophic clones amongst the infected Jurkat population and thus proceeded to selection for valine prototrophic clones.

Jurkat-Div cells were cultured on low valine medium (0 nM; 1.7 nM; 4.25 nM) alongside control cells infected only with GFP (Jurkat-GFP) (**Extended Data 8A**). At all valine concentrations, both cell populations saw a reduction in cell number over 14 days. By day 21, however, Jurkat-Div cells saw an increase in cell number across all conditions while Jurkat-GFP cells had either died off or remained at low cell counts. We serially diluted the resulting selected Jurkat-Div cell populations (Jurkat-Val) to isolate and expand 119 individual clones in valine-replete conditions prior to characterization via amino acid dropout assay and Amplion-seq (**Figure 5A**). As before, we considered a clone prototrophic if its Val Score was at least 2X that of a parental control (Val Score > 0.23), and 71% (85/119) of clones met this threshold (**Figure 5B**). We further characterized two clones exhibiting valine prototrophy, Jurkat-Val-31 and Jurkat-Val-32, culturing both clones in valine-free medium over 11 days and measuring their absolute growth rate. While control cells exhibited stagnant cell numbers over 11 days of culture, clones Jurkat-Val-31 and Jurkat-Val-32 proliferated at 2.02 and 1.87 days/doubling, respectively, corresponding to 50% and 52% of their growth rates in valine-replete medium (**Figure 5C; Extended Data 8B**).

**Figure 5.**
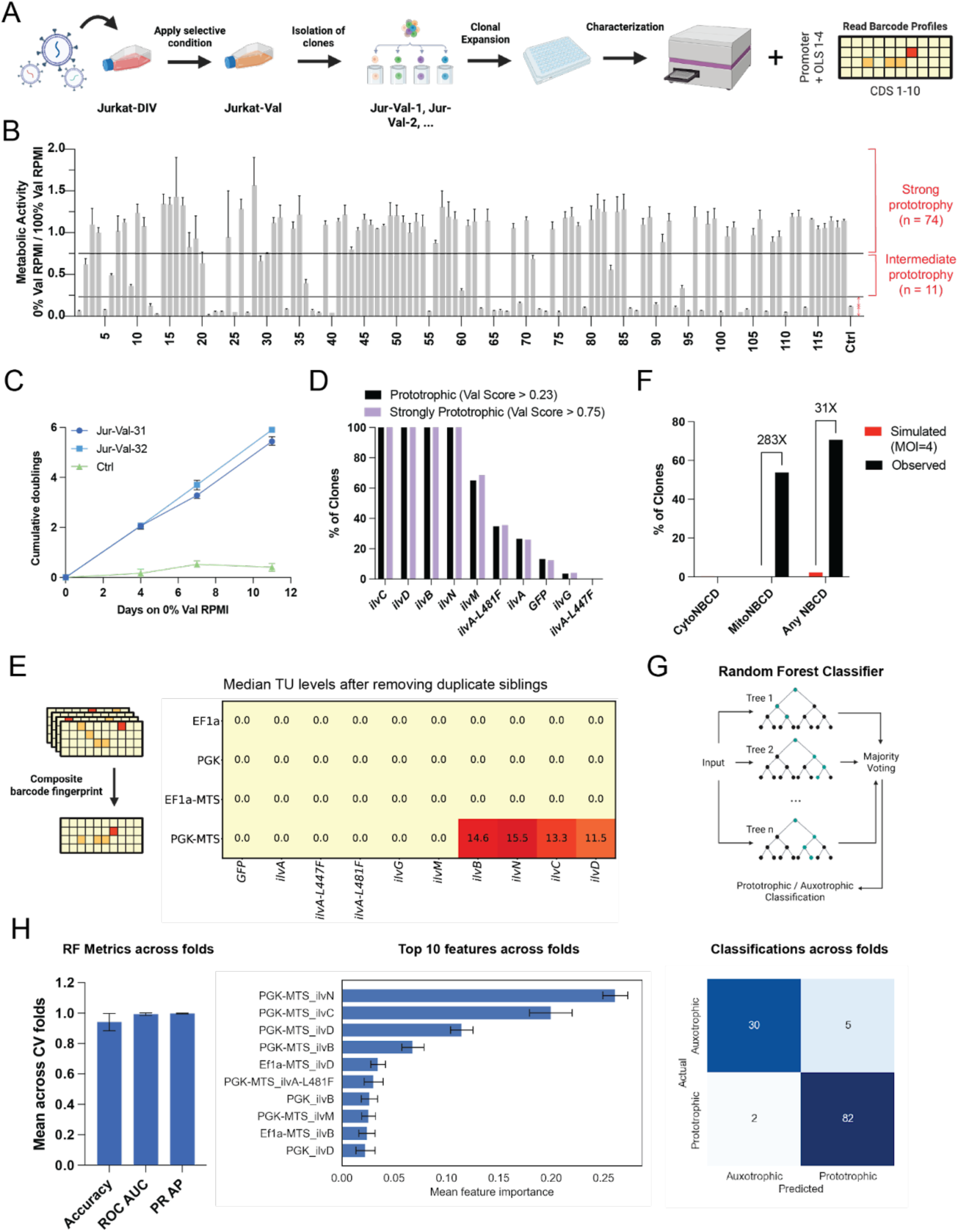
**Discovery of a functional valine biosynthesis pathway in Jurkat cells** A. Schematic detailing SGE workflow from infection to phenotyping and barcode readout for suspension cells. B. PrestoBlue amino acid dropout assay of 119 individual clones selected on low valine RPMI. Each clone was measured in triplicate wells in valine-free medium for 3 days and relativized to triplicate wells with valine-replete medium for 3 days to generate a ‘Val Score’. Valine prototrophy was defined as a 2X improvement in Val Score relative to a non-prototrophic parental cell line. Strong valine prototrophy was defined as a Val Score of 0.75. Error bars represent standard deviation across triplicate wells. C. Growth curve of Clones Jur-Val-31 and Jur-Val-32 cultured in valine-free RPMI and compared to a non-prototrophic parental cell line. Error bars represent standard deviation across triplicate wells. D. Monitoring CDS presence across all valine prototrophic clones (Val Score > 0.28) and across all strongly valine prototrophic clones (Val Score > 0.75). E. Composite barcode ‘fingerprint’ comprised by median normalized read count across all strongly valine prototrophic clones (Val Score > 0.75) after removing duplicate ‘sibling’ clones. F. Comparing representation of *ilvN, ilvB, ilvC,* and *ilvD* in clones selected on low valine medium relative to their representation if selected by random chance as determined by a simulated infection at MOI=4. G. The Random Forest model classifies outcomes using a majority vote across many decision trees trained on subsets of data and features. H. 4-fold cross-validation was used to ensure stability of the RF model. Metrics (left), top 10 features (middle), and classifications (right) across the 4 folds are summarized. Error bars represent the standard deviation across the 4 cross-validation folds.

To discover the underlying combinations of TUs that comprised valine prototrophy solutions in Jurkat, we again performed amplicon-seq on isolated clones. Clones Jurkat-Val-31 and Jurkat-Val-32 exhibited similar TU profiles with mitochondrially localized *ilvM, ilvB, ilvN, ilvC,* and *ilvD* in their PGK promoter context as well as cytoplasmically localized *ilvB* in its PGK promoter context (**Extended Data 8C**). Both clones appeared to carry 9 TU integrations corresponding to 32 kb of integrated DNA of which 15 kb was directly made up by TU modules. We then identified the CDSs universally represented amongst clones exhibiting valine prototrophy by setting a Z-score threshold of 1.15 as before (**Extended Data 8D**). *ilvB, ilvC, ilvD*, and *ilvN* were present in all Jurkat valine prototrophic clones, matching the CDS content of the solution previously identified for the CHO-Div population (**Figure 5D; Extended Data 8E**). In order to ascertain the optimal promoter and localization context for each CDS, we again generated a composite barcode ‘fingerprint’ consisting of the median representation measured for each TU in the library. In doing so we found *ilvC, ilvD* and *ilvN* were favored in their mitochondrially localized context under control of the PGK promoter while ilvB appeared to be favored under control of a PGK promoter in both the cytoplasm and mitochondria (**Extended Data 8F**). However, when looking at the 3 promoter barcodes and 3 CDS barcodes underlying these 5 TUs, we noticed that many clones exhibited barcode signatures shared with other clones suggesting that multiple sequenced samples had arisen from the same founder clone (**Extended Data 8G**). We expect to observe these clonal ‘siblings’ if a given pathway solution is especially outperforming others in the population and a selective sweep of the population occurs. This prompted us to look more closely at samples with globally correlated TU signatures. We found that 69 clones had at least 1 ‘sibling’ in the population while 50 clones were unique ‘singletons’ (**Extended Data 9A**). Using graph-based clustering, we found 10 separate sibling groups varying in size from 2-24 clones, and sibling group size did not correlate to average Val Score (**Extended Data 9B**). We examined the 3 biggest sibling groups - 2, 4, 9, and 5 - consisting of 24, 12, 10, and 7 clones, respectively, and looked at representative heatmaps for each group (**Extended Data 9C**). A unifying theme across the three sibling groups was mitochondrial localization of *ilvN, ilvC*, and *ilvD* with *ilvB* being localized to either the cytoplasm, mitochondria or to both compartments. We next removed duplicate siblings to generate a composite barcode fingerprint of all strongly prototrophic samples amongst unique clones only, which produced pathway solution consisting only of mitochondrially localized *ilvN, ilvB, ilvC*, and *ilvD* (**Figure 5E**). When looking at how overrepresented this pathway solution was compared to simulated expectation, we found the collective grouping of *ilvN, ilvB, ilvC*, and *ilvD*, irrespective of subcellular compartmentalization, was overrepresented 31X while the collective mitochondrial localization of these CDSs was overrepresented 283X (**Figure 5F**).

We also looked at CDS representation amongst 24 valine auxotrophic Jurkat clones carrying at least 1 integration (**Extended Data 10A**). Unlike the auxotrophic CHO clones selected on low isoleucine medium, which all carried a partial isoleucine biosynthetic pathway, selection for partial biosynthetic pathways in Jurkat clones selected on low valine medium was much less pronounced with slight CDS overrepresentation of *ilvD, ilvC,* and *ilvB* relative to expectation (**Extended Data 10B)** perhaps a reflection of the lower MOI used to infect Jurkat cells or perhaps indicating that clonal resource-sharing is less pronounced in suspension cell culture.

### Jurkat-Div cells selected on low isoleucine medium

We also subjected Jurkat-Div cells to a range of low isoleucine concentrations (0 nM; 6 nM; 12 nM Ile) in the event that we might discover a functional isoleucine biosynthetic pathway in Jurkat cells. Across the 3 conditions and 32 days of culture, proliferation of Jurkat-Div cells was never found to outpace that of Jurkat-GFP control cells suggesting that no isoleucine prototrophic phenotypes were present in the population (**Extended Data 11**). This is not surprising given that we simulated the presence of TUs that made up the strongly isoleucine prototrophic phenotypes seen in CHO and found little representation of these combinations in Jurkat-Div cells at MOI=4 relative to CHO-Div cells at MOI=8 (**Extended Data 7; Figure 4K**). This indicates that simulating pathway diversity based on estimated TU distributions and MOI can effectively inform pathway engineering choices.

### Machine Learning to Predict Prototrophy Phenotypes from SGE-generated data

Finally, we asked whether machine learning could be used to predict prototrophy phenotypes directly from the TU composition of each clone. We decided to apply a random forest model for classification due to its strength in interpreting non-linear relationships between features, its robustness to overfitting, and its ability to provide feature importance scores (**Figure 5G**). Using a Val Score threshold of 0.23, samples were binarized into prototrophic or auxotrophic. The samples were partitioned into four mutually-exclusive class-balanced folds to perform stratified 4-fold cross-validation (**Extended Data 12**). For each fold, the model was trained on the other three folds, and evaluated on the held-out fold for 4 separate 75:25 train:test splits. When looking at metrics across folds, the model successfully classified samples into prototrophic or auxotrophic with a mean accuracy of 0.942 +/- 0.005, a mean ROC AUC of 0.993 +/- 0.006 and a mean precision-recall AUC of 0.997 +/- 0.002 across folds, indicating strong discriminative ability and stability across different train:test sample compositions (**Figure 5H**). In total, 112/119 clones were correctly classified, and the top 4 features driving classification were mitochondrially localized *ilvN, ilvC, ilvD* and *ilvB* under control of a PGK promoter, matching the pathway solutions that we discovered by manual analysis and interpretation. These results demonstrate that SGE-generated data are well-suited for machine learning-based modeling and that such models can effectively extract biologically meaningful features, enabling pathway-scale analysis for solutions to complex metabolic engineering problems.

## DISCUSSION

Here, we introduce Shotgun Genetic Engineering, a method for efficiently exploring vast combinatorial design spaces comprising multiple interacting transcriptional units (TUs) in mammalian cells. By leveraging single cells as independent experiments and using random high-copy integration of individual TUs per cell, SGE enables high-throughput assembly and screening of millions of synthetic metabolic pathways. This approach bypasses the inefficiencies associated with building and delivering large DNA constructs, substantially accelerating pathway discovery and optimization.

Using this approach, we generated and screened millions of synthetic metabolic pathways to engineer essential amino acid prototrophy in mammalian cells. We achieved

i. near wild-type growth in valine-free medium, and (ii) the first demonstration of isoleucine prototrophy in mammalian cells. The functional solutions involved integration of 23-52 kb of synthetic DNA - construct sizes impractical to screen using conventional methods. We also provide initial evidence that leveraging organellar compartmentalization can enhance engineered pathway functionality. Finally, we demonstrate the versatility of SGE across mammalian cell types (species, suspension/adhesion) and present a pilot integration with machine learning to predict functional pathway combinations.

A consistent theme across the metabolic engineering efforts described here was the preference for mitochondrial localization of BCAA pathway enzymes. This was unexpected given that the genes in our library were sourced from *E. coli*, a prokaryote lacking subcellular compartments. However, mitochondria are thought to be of bacterial origin and share significant protein composition with bacteria, which may help explain this preference^28^. Interestingly, equivalent BCAA biosynthesis genes in extant eukaryotic BCAA prototrophic microbes such as fungi are also localized to mitochondria providing a clue that there may be biochemical advantages associated with mitochondrial localization such as increased pyruvate concentrations relative to cytosol^29,30^ or greater availability of relevant pathway cofactors e.g. iron-sulfur clusters and thiamine diphosphate^31–33^. Similarly, in plants, these BCAA biosynthetic enzymes are targeted to chloroplasts which also are an abundant source of pyruvate^34^. Another possibility is that BCAA biosynthetic enzymes simply benefit from being much more concentrated in a limited organellar physical space, thereby boosting pathway flux and function as has previously been demonstrated for other pathways localized to the mitochondria in *S. cerevisiae*^35^. This finding highlights the utility of incorporating organellar diversity in the SGE screen, and encourages incorporation of yet more organellar targeting signals in future engineering efforts in mammalian cells.

Use of SGE comes with several limitations, most notably the requirement for a selectable phenotype. Essential nutrients provide a built-in live/death screen that effectively identifies cells carrying optimal solutions. However, when the desired phenotype is not linked to cell survival, alternative selection strategies such as coupling pathway activity to fluorescent biosensors, surface marker expression, or engineered genetic circuits that connect pathway output to a selectable reporter may be applied^36,37^. Another limitation of SGE as currently implemented is the use of lentivirus for random integration. We observed significant biases in TU representation introduced during viral packaging. Specifically, EF1a-driven constructs were markedly underrepresented compared to those with PGK promoters, and similar biases were noted for certain coding sequences in ways not attributable solely to size. Transitioning to transposon-based delivery systems could alleviate this issue. Finally, while amplicon-seq can inform what TUs have been integrated in clones of interest, it provides little information about which genes are expressed. In cell lines prone to silencing, this may confound interpretation. Pairing SGE with either bulk or single-cell RNA-seq would provide expression-level information, enabling more accurate assessment of functional pathway configurations.

While SGE excels at rapidly exploring vast design spaces via random integration, successful multi-TU solutions identified in this manner may need to be reconstituted into compact, single-vector formats for practical deployment. However, transcriptional units may behave differently when assembled into condensed architectures^38^. A hybrid strategy may be effective in this case - using SGE for high-throughput discovery, followed by CLASSIC^11^ or related methods for validation and optimization in compact constructs in lower throughput.

Despite its current limitations, SGE represents a powerful strategy for the scalable engineering of increasingly complex biosynthetic traits in mammalian cells. In this study, we demonstrated the discovery of a functional six-gene isoleucine biosynthesis pathway from a 40-member TU library. We anticipate that SGE can be extended to larger libraries by proportionally increasing the MOI during integration, enabling access to even more complex phenotypes. These may include prototrophy for additional essential amino acids, growth factors, and vitamins; enhanced tolerance to environmental stressors such as hypoxia, shear stress, and metabolic byproducts; or rewiring of metabolic programs toward alternative cell states or identities. The inclusion of elements that modulate host factors (CRISPRa/i machinery) would further expand the design space explored by this approach.

Ultimately, the ability to engineer mammalian cells with complex pathways has broad implications across basic research, biotechnology, and medicine. For instance, a mammalian cell engineered to be prototrophic for all amino acids would require a reduced number of nitrogen sources, opening the door to utilizing stable isotope labeling to chart presently unknown corners of the mammalian metabolic map^39^. Meanwhile, mammalian cells engineered to become independent of specific nutrients could lower media costs and simplify formulations for biomanufacturing applications such as cultivated meat and viral vector production^7,40^. Finally, cell therapies may be metabolically augmented to better withstand the hostile tumor microenvironment, which is marked by nutrient deprivation, acidosis, oxidative stress and inflammation with potential for improving the performance of cell-based therapeutics used in the treatment of solid tumors^41–44^.

## METHODS

### Cell lines, Media, and Cell Counting

CHO Flp-In cells (ThermoFisher, R75807) and Jurkat cells (M. Pacold Lab, NYU) were used in all experiments. All cell lines tested negative for mycoplasma. For growth assays involving amino acid dropout formulations, medium was prepared from an amino acid-free RPMI 1640 powder base (US Biological, R8999-20), and custom combinations of amino acids were added back in as needed to match the standard amino acid concentrations for RPMI 1640 except for isoleucine, which was considered 100% supplemented at 0.06 mM. Additionally, RPMI medium was supplemented with 2 mM sodium pyruvate as previously described^18^. Custom amino acid dropout medium was adjusted to a pH of 7.3 with HCl, sterile filtered, and supplemented with 10% dialyzed Fetal Bovine Serum (Fisher Scientific, 26400044) and Penicillin-Streptomycin (100 U/mL) prior to use. For metabolomics experiments, medium was prepared from an amino acid-free, glucose-free RPMI 1640 powder base (US Biological, R9010-01) and custom combinations of amino acids were added in to match the standard concentrations for RPMI 1640. In the place of glucose, isotopically heavy ^13^C_U_ glucose (Cambridge Isotope Labs, CLM-1396-PK) and ^13^C_3_ sodium pyruvate (Millipore Sigma, 490717) were added at 11.1 mM and 2 mM, respectively. Cell counts were performed using a Countess 3 FL.

### SGE Library Construction

Promoters and OLSs were PCR amplified and allotted unique barcodes by use of PCR primers encompassing said barcodes and BsmBI restriction sites. Each barcoded promoter/OLS combination was cloned into an assembly vector with lentiviral long terminal repeats using Gibson assembly. CDSs of interest were codon optimized and synthesized commercially. Each CDS was allotted 3 unique barcodes by PCR amplification with primers encompassing said barcodes along with BsmBI restriction sites to facilitate downstream golden gate cloning. Each PCR-amplified, barcoded CDS was cloned into a pCR-Blunt-II-TOPO vector. Concentrations of each TOPO-cloned CDS were measured using a Qubit fluorometer prior to pooling of all CDSs of interest at equimolar concentration. Concentrations of each promoter/OLS plasmid were similarly measured using a Qubit fluorometer prior to pooling of all promoter/OLS plasmids at equimolar concentration. The pooled CDS mixture was spiked into the golden gate reaction at 5X the molar concentration of the pooled assembly vector mix, which was used at 100 ng. We added 1.5 ul 10X NEB T4 ligase buffer, 0.15 ul BSA, 1.1 ul BsmBI-v2 (10,000 U/mL), 1 ul T4 ligase (20,000 U/mL) and then supplemented with H_2_O to a 15 ul total volume. Reactions were run for 90X cycles of 42°C, 3 min and 16°C, 4 min prior to 1 cycle of 50°C, 5min and 80°C, 5min. Resulting libraries were transformed into ElectroMAX Stbl4 *E. coli* (Invitrogen, 11635-018) for amplification prior to library extraction and amplicon-seq to measure library distributions.

### Lentiviral packaging and infection

Lentivirus was packaged by plating 4×10^6^ HEK293T cells on 10 cm^2^ plates and incubating cells overnight at 37°C. Cells were transfected with a plasmid mix consisting of 3.5 µg of the SGE library, 6.0 µg psPAX2 (Addgene, 12260), and 3.0 µg pMD2.G (Addgene, 12259) using Lipofectamine 2000 (ThermoFisher Scientific, 11668019) in accordance with the manufacturer’s instructions. Transfected HEK293T cells were incubated for 48 hrs, before medium was collected and replenished. Cells were then incubated for an additional 24 hrs before medium was collected again. Collected medium was centrifuged at 200xg for 5 mins, and the resulting supernatant was filtered using a 0.45 µm filter. Virus was concentrated using Milipore Amicon Ultra-15 Centrifugal filter units (MiliporeSigma, UFC903024) for 30 mins at 4000xg prior to storage at −80C until use. For CHO infections, cells were plated at 1.5M cells per 10cm^2^ and incubated overnight prior to infection. At time of infection, medium was replaced with fresh medium containing polybrene at 8 µg/mL and concentrated virus. Cells were incubated with the virus for 24 h before replenishing with fresh medium. For Jurkat infections, 3M cells were resuspended in 6 mL medium containing polybrene at 8 ug/mL and concentrated virus prior to centrifugation at 931xg for 2 h at 30°C. An additional 6 mL of medium (not containing polybrene) was subsequently added to cells dropwise prior to 24 h incubation at 37°C after which medium/virus was replenished with fresh medium.

### MOI estimation by qPCR

gDNA was extracted from cells using the QIAamp DNA Mini Kit (Qiagen, 51304) in accordance with the manufacturer’s protocol. qPCR reactions were performed as 10 ul reactions with 0.5 µl 5 µM primer mix and using SYBR Green Master I on a QuantStudio 7 Pro. Primers were designed to amplify amplicons 60-120 bp in size. MOI was estimated by measuring amplification of a sequence universally found in each TU found in the SGE library and relativizing to amplification of CHO targets, Dhfr & Hprt (expected to be single copy), and Jurkat targets, RPP30 & ALB (expected to be found in two copies), respectively, using the ΔCT method. All reactions were performed in triplicate.

### Monte Carlo simulation of TU integrations across variable MOIs

To model the distribution of transcription unit (TU) combinations integrated into a cell population, we performed Monte Carlo simulations assuming Poisson-distributed multiplicities of infection (MOI). For CHO cells, we simulated 2,420,000 cells at MOI values of 2, 4, 6, 8, 10, 12, and 14, spanning a range around the estimated MOI of 8.8. For Jurkat cells, 3,000,000 cells were simulated at MOIs of 2, 4, 6, and 8, centered around the estimated MOI of 3.4. Integration events were simulated by first drawing the number of integrations per cell from a Poisson distribution. For cells with one or more integrations, TUs were sampled with replacement according to empirically defined distributions. The resulting sets of integrated TUs were sorted and recorded as combinations, irrespective of integration order. For each MOI, the frequency of unique TU combinations was tallied and used to benchmark the probability of observing specific TU sets.

### Amino Acid Prototrophy Screens

We screened for amino acid prototrophic clones using PrestoBlue HS Cell Viability reagent (ThermoFisher, P50200), a resazurin-based viability reagent that fluoresces in response to cellular reducing activity. Fluorescence intensity is proportional to the number of metabolically active (viable) cells in the medium. For each amino acid dropout condition, we relativized triplicate wells for each clone in the amino acid dropout condition to triplicate wells in an amino acid replete condition to control for differences in overall growth rate and seeding density. Amino acid replete medium was made by making amino acid dropout medium and adding in the missing amino acid. Fluorescence measurements were conducted on an Agilent Biotek Synergy Neo2.

### Amplicon-Seq Library Preparation

gDNA from clones of interest was prepared using the Zymo Quick-DNA 96 Plus kit (Zymo, D4070) in accordance with the manufacturer’s protocol. PCR1 (UMI addition) was performed using primers annealing to a universal primer sequence found in the 3’ UTR across all SGE TU vectors. PCR 1 used 250 ng gDNA template, 2.5 µl of each 10 µM primer and Q5 polymerase in 50 µl total reaction volume. PCR1 cycling conditions were 98°C - 5 min; 4 cycles of 98°C - 20s, 65°C - 20s, 72°C - 30s; 72°C - 60s. PCR clean up was performed using AmpureXP beads (0.8X) and eluted in 15 µl 0.1X TE. For PCR2 (sample indexing and sequencing adapter addition), 10 µl of eluate from the previous reaction was used in another 50 µl reaction using Q5 polymerase and 5 µl of a 5 µM indexing primer mix. SybrGreen dye was also added to the reaction mix to allow us to monitor the reaction progression and stop reactions prior to saturation (typically 16-19 cycles). PCR2 cycling conditions were 98°C - 5 min; 16 cycles of 98°C - 10s, 65°C - 10s, 72°C - 20s. PCR clean up was again performed using AmpureXP beads (0.8X) and eluted in 15 µl 0.1X TE. Sequencing was performed on a NextSeq500 75 cycle kit to yield PE 36,36 reads.

### Amplicon-Seq Analysis

FASTQ files from paired ends were merged using pear (v0.9.11). Stitched FASTQ files were processed to recover known barcode combinations identifying promoter/OLS and CDSs, respectively, to thus quantify transcription unit (TU) integration events. Reads were parsed to identify two universal priming regions flanking the barcode segments. For reads containing both universal primers, two 8-bp barcodes were extracted and concatenated to form a unique identifier for each TU.

Each extracted barcode combination was matched against a known barcode reference library for exact matches. However, we noticed that a subset of clones exhibited a large proportion of reads that did not match any in the reference library but exhibited a 1-bp mismatch relative to reads that were found in the reference library. Since no two barcodes in the library exhibited a Hamming distance <2, and 99.9% of possible barcode pairings exhibited a Hamming distance of 3 or more, we opted to allow for 1-bp mismatches if possible to do so unambiguously. Briefly, if no exact barcode match was found, sequences were compared to a precomputed lookup table containing all possible 1-bp mismatch variants of the reference barcodes. Reads with a unique (unambiguous) 1-bp mismatch were assigned to the corresponding reference barcode, while ambiguous 1-bp mismatches (matching multiple barcodes) were excluded from downstream quantification. Samples with fewer than 50,000 matching reads were further excluded from downstream analysis.

For analysis of CDS presence within each clone, read counts for each TU were used to calculate a Z-score. Given 3 barcodes per promoter/OLS combination, and 3 barcodes per CDS, each TU has 9 underlying barcodes. Read counts for each TU were aggregated and log-transformed prior to calculating a Z-score for each TU on a per sample basis to separate true signal from background. For generating heatmaps to identify the optimal combinations of TU amongst subsets of clones, read counts were normalized to 1 to account for sequencing depth prior to finding the median.

### Metabolomics

Cells were cultured in RPMI medium containing ^13^C-glucose and ^13^C-sodium pyruvate with or without valine/isoleucine (as indicated) prior to cell harvest. Cell pellets were generated by trypsinization, followed by low-speed centrifugation, and the pellet was frozen at –80°C until further processing. A metabolite extraction was carried out on each sample with an extraction ratio of 1e6 cells per mL (80% methanol containing internal standards, 500 nM). The LC column was a Millipore ZIC-pHILIC (2.1×150 mm, 5 μm) coupled to a Dionex Ultimate 3000 system and the column oven temperature was set to 25°C for the gradient elution. A flow rate of 100 μL/min was used with the following buffers; (A) 10 mM ammonium carbonate in water, pH 9.0, and (B) neat acetonitrile. The gradient profile was as follows; 80–20%B (0–30 min), 20–80%B (30–31 min), 80–80%B (31–42 min). Injection volume was set to 1 μL for all analyses (42 m total run time per injection).

MS analyses were carried out by coupling the LC system to a Thermo Q Exactive HF mass spectrometer operating in heated electrospray ionization mode (HESI). Method duration was 30 min with a polarity switching data-dependent Top 3 method for both positive and negative modes, and targeted MS2 scans for the monoisotopic, U-13C, and U-13C/U-15N valine *m*/*z* values. Spray voltage for both positive and negative modes was 3.5 kV and capillary temperature was set to 320°C with a sheath gas rate of 35, aux gas of 10, and max spray current of 100 μA. The full MS scan for both polarities utilized 120,000 resolution with an AGC target of 3e6 and a maximum IT of 100 ms, and the scan range was from 67 to 1000 *m*/*z*. Tandem MS spectra for both positive and negative mode used a resolution of 15,000, AGC target of 1e5, maximum IT of 50ms, isolation window of 0.4 m/z, isolation offset of 0.1 m/z, fixed first mass of 50 m/z, and 3-way multiplexed normalized collision energies (nCE) of 10, 35, 80. The minimum AGC target was 1e4 with an intensity threshold of 2e5. All data was acquired in profile mode. All valine/isoleucine data was processed using Thermo XCalibur Qualbrowser for manual inspection and annotation of the resulting spectra and peak heights referring to authentic valine/isoleucine standards and labeled internal standards as described.

### Identification and Analysis of ‘Sibling’ Clones

To detect samples with globally similar barcode profiles, all pairwise comparisons between samples were computed using a Pearson correlation coefficient. Pairs were retained if each sample had at least two non-zero barcode features (to avoid spurious correlations from sparse data), and the Pearson correlation coefficient between the sample profiles was ≥ 0.9. To organize putative sibling clones identified through barcode profile correlation into groups, we applied a graph-based clustering approach. The set of high-correlation sample pairs was used to construct an undirected graph in which each node represented a sample and each edge represented a strong global match using the NetworkX Python package with edges connecting sample pairs that met the correlation threshold. Connected components within the graph were then identified, each representing a cluster of samples inferred to be derived from the same original clone or barcoded integration event. Samples not belonging to any component were defined as singletons.

### Random Forest Classification

To identify transcription unit (TU) features predictive of valine prototrophy, we trained a Random Forest classifier using normalized read counts as input features. A binary phenotype was defined by thresholding the Val Score at 0.23, classifying samples as either prototrophic or non-prototrophic. We used stratified 4-fold cross-validation to ensure balanced class distributions across splits. In each fold, a Random Forest classifier was trained on 75% of the data and evaluated on the remaining 25%. Model performance was assessed using confusion matrices, classification reports, receiver operating characteristic (ROC) curves with area under the curve (AUC), and precision-recall (PR) curves with average precision (AP). Confusion matrices were plotted per fold, and mean performance metrics were reported across folds with standard deviation. Feature importances were extracted from each trained model to identify the most predictive TUs. The top 10 features were reported per fold, and mean feature importances were calculated to identify features consistently ranked as important across folds.

## ACKNOWLEDGEMENTS

We thank M. Pacold for sharing Jurkat cells and members of the Boeke Lab for helpful discussions. Figure schematics were made using <u>Biorender.com</u>, and ChatGPT provided assistance in generating code for data analysis. We thank M. Maurano and his team for help with DNA sequencing. Supported in part by NHGRI/NIH grant RM1-HG009491 to JDB, NIH grant DP5OD036167 to SP and by grant 2024-349901 from the Chan Zuckerberg Initiative to SP.

## AUTHOR CONTRIBUTIONS

JT and SP conceptualized the project. JT, SS, and TR performed experiments. JT developed lab automation pipelines, computational pipelines and performed data analysis. DRJ supervised metabolomics experiments. DF supervised mathematical modeling. SP and JDB supervised all other work. JT, SP, and JDB wrote the manuscript with input from all authors. JDB secured funding for the project.

## DATA AVAILABILITY

All scripts written to analyze data and generate figures can be accessed at https://github.com/julietrolle/shotgun-genetic-engineering. Raw data was deposited to the sequencing read archive (SRA) under PRJNA1282762.

---

**Extended Data 1.**
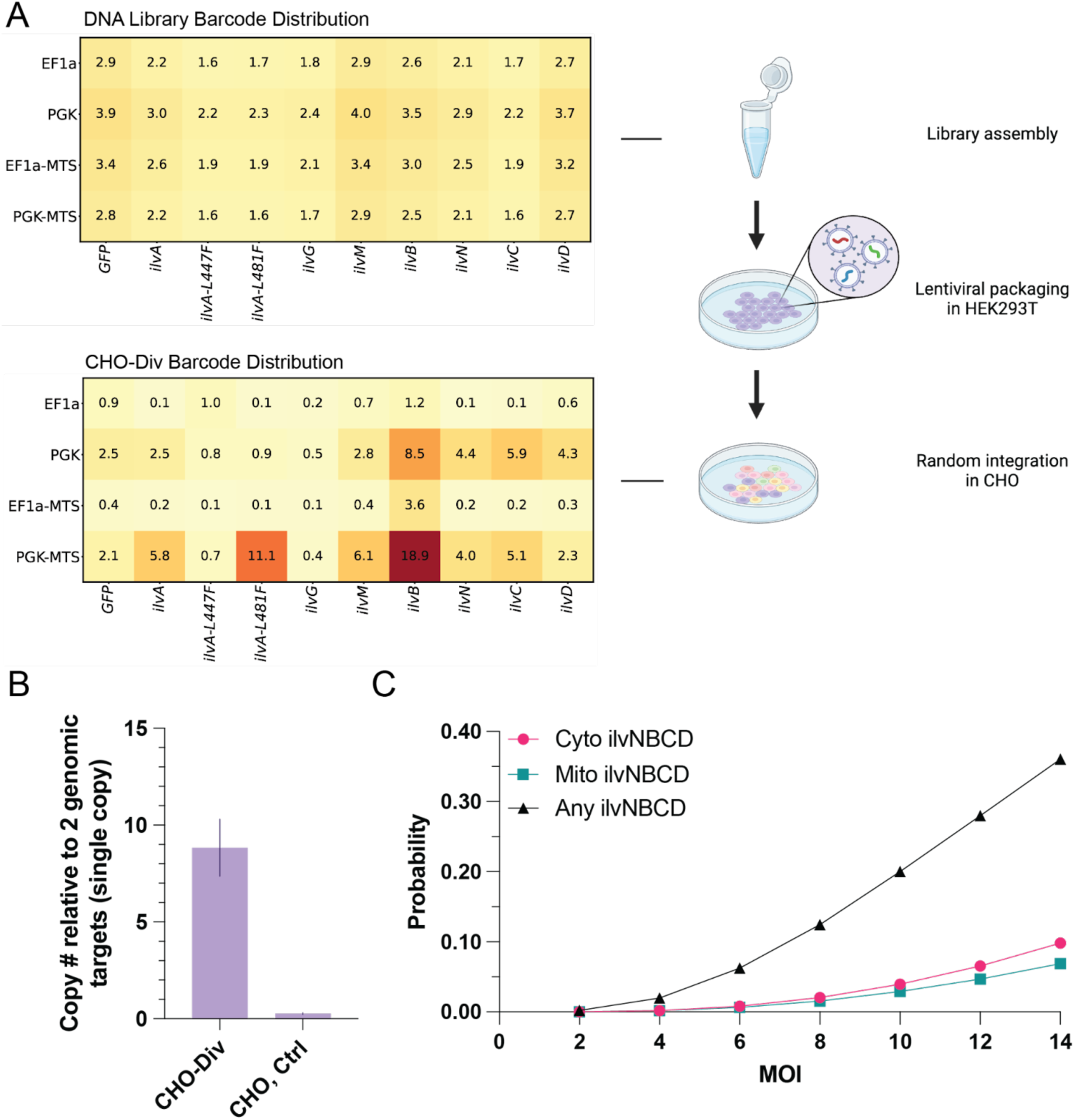
**Transcription unit distribution in CHO pre-infection (DNA Library) and post-infection (CHO-Div) of SGE library.** A. Transcription unit distribution prior to lentiviral packaging compared to transcription unit distribution after infection of CHO cells. Y-axis indicates promoter/OLS context while the X-axis indicates CDS. Numbers are percentages i.e. all TU numbers add up to 100%. B. qPCR to determine copy number of TU integrants relative to two single copy genomic targets (Dhfr, Hprt) for estimation of MOI. C. Probability of finding a known valine biosynthesis pathway combination in CHO-Div based on a Monte Carlo simulated infection of 2.42M cells with the library distribution measured for CHO-Div cells in (A). Depicted is the probability of finding *ilvN, ilvB, ilvC* and *ilvD* irrespective of subcellular compartmentalization (black), all 4 genes localized to the cytoplasm (red), or all 4 genes localized to mitochondria (green) at a range of MOIs.

**Extended Data 2.**
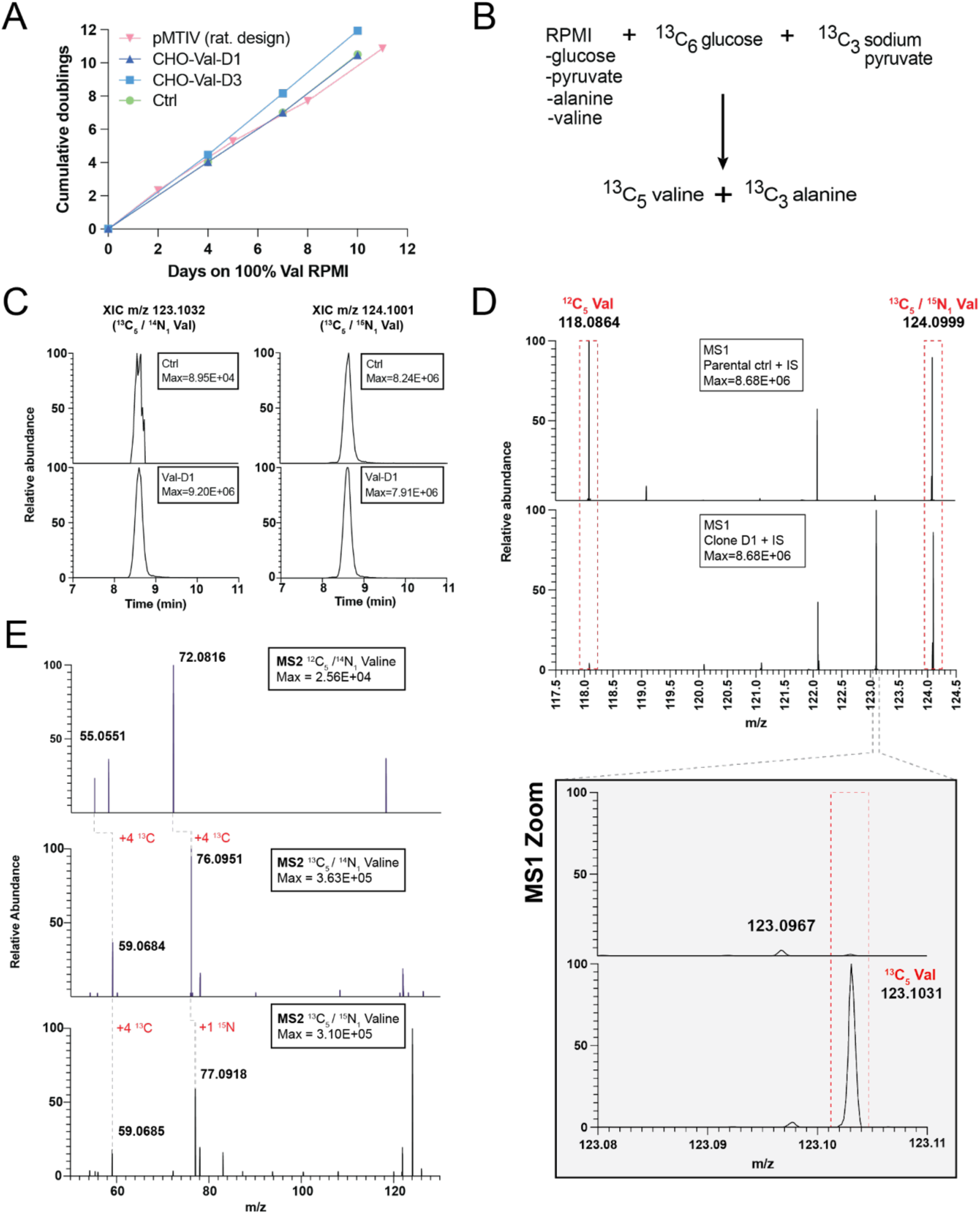
**Characterization of CHO-Div clones selected on low valine medium, including ^13^C tracing to confirm endogenous biosynthesis of valine in Clone CHO-Val-D1.** A. Growth curve of Clones CHO-Val-D1 and CHO-Val-D3 cultured in valine-replete RPMI compared to the parental cell line. Error bars represent standard deviation across triplicate wells. B. Schematic detailing ^13^C labeling strategy to detect biosynthesized valine. Prototrophic cells were cultured on valine-free ^13^C labeled medium while control cells were cultured in a ^13^C valine-replete equivalent. C. Extracted ion chromatography shows detection of ^13^C Valine in Clone CHO-Val-D1, which matches the retention time of spiked-in internal standard ^13^C_5_/^15^N_1_ Valine. Detection of ^13^C Valine in parental control cells is minimal by comparison. D. MS1 peak corresponding to ^13^C_5_ Valine is seen for Clone CHO-Val-D1 and not for the parental control cells E. MS2 confirms that fragmentation patterns for putative ^12^C_5_ valine, ^13^C_5_ valine and ^13^C_5_/^15^N_1_ valine match expectation.

**Extended Data 3.**
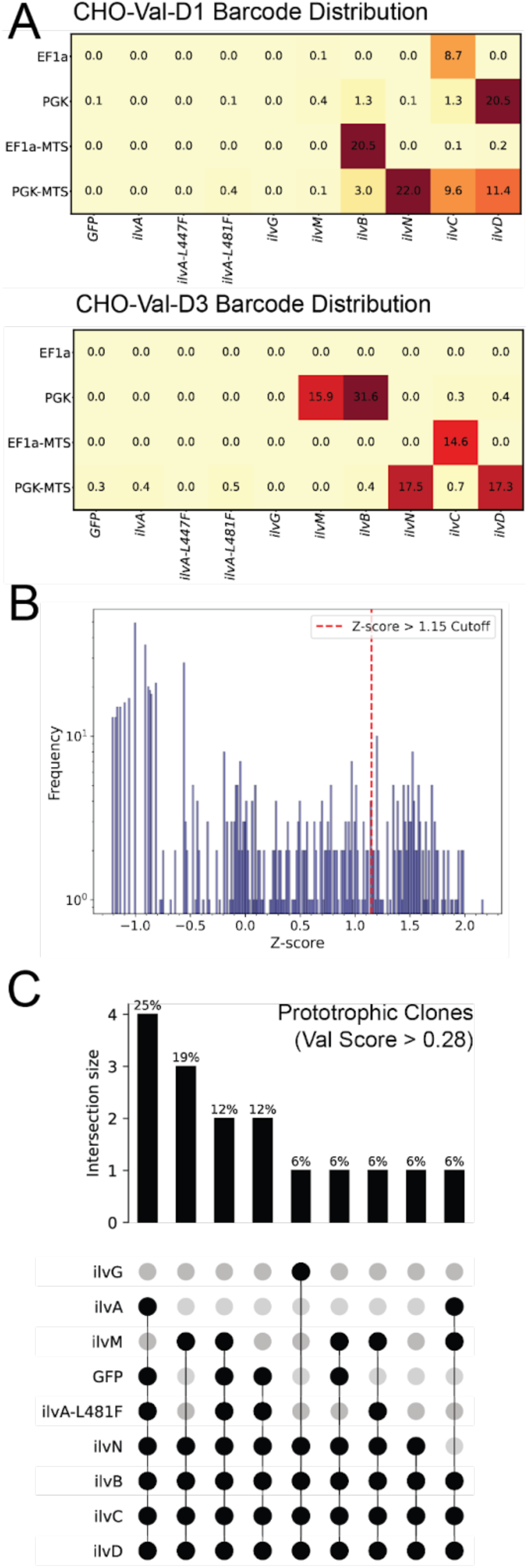
**Coding sequence usage underlying sequenced clones selected on low valine RPMI.** A. Normalized read counts for each TU, Clones CHO-Val-D1 and CHO-Val-D3. B. Log-transformed read counts were used to calculate a Z-score for each TU in each sample. A Z-score threshold of 1.15 was used to define TU presence vs. TU absence across the 16 clones. C. An UpsetPlot depicting which combinations of CDSes were determined to be present within each of the 16 valine prototrophic clones.

**Extended Data 4.**
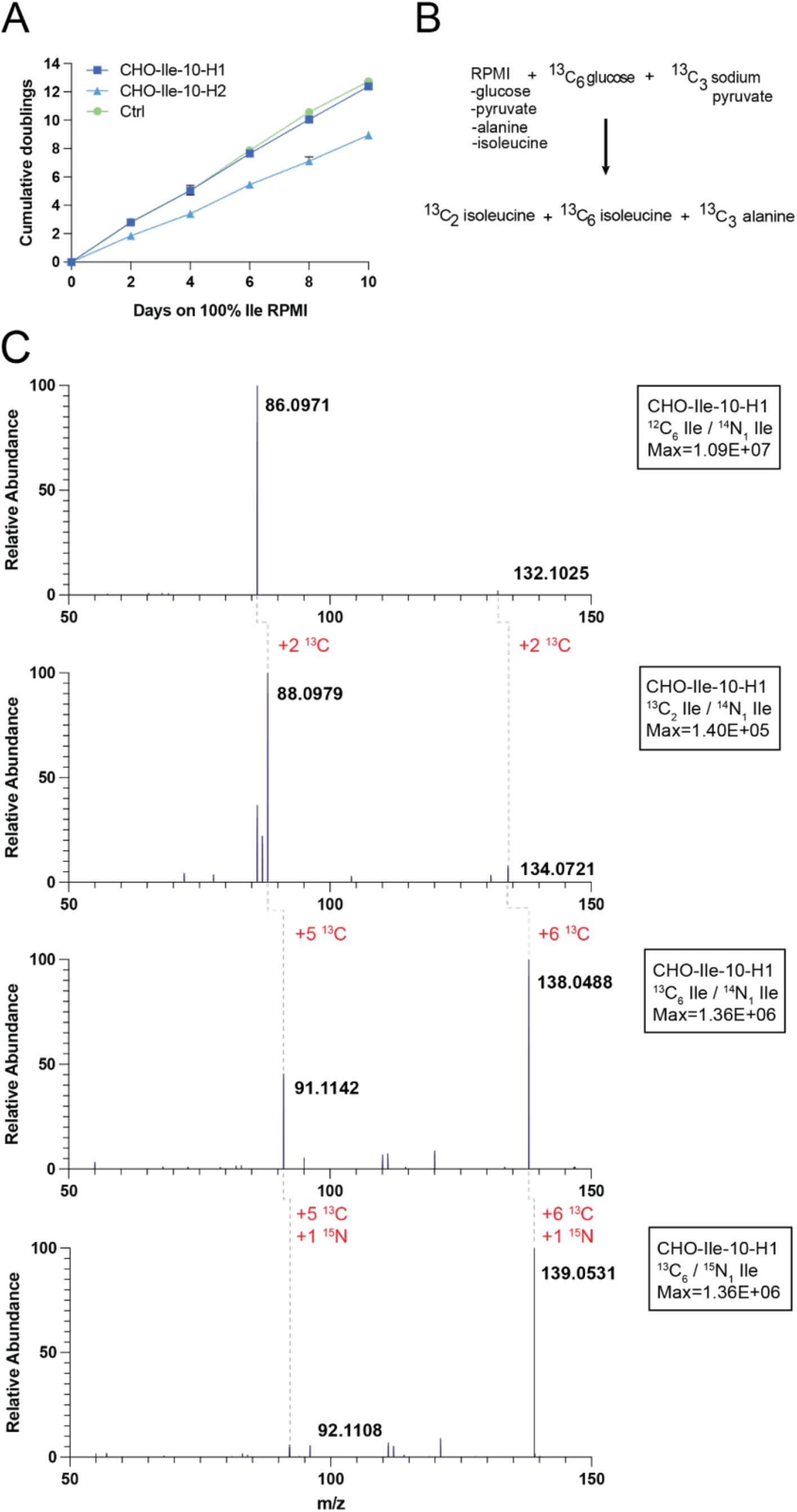
**Characterization of CHO-Div clones selected on low isoleucine medium including ^13^C tracing to confirm endogenous biosynthesis of isoleucine in Clone CHO-Ile-10-H1.** A. Absolute cell counts of clones CHO-Ile-10-H1 and CHO-Ile-10-H2 cultured on isoleucine-replete medium compared to the parental cell line. Error bars represent standard deviation across triplicate wells. B. Schematic detailing ^13^C labeling strategy to detect biosynthesized isoleucine. Prototrophic cells were cultured on isoleucine-free ^13^C labeled medium while control cells were cultured in a ^13^C isoleucine-replete equivalent. C. MS2 confirms that fragmentation patterns for putative ^12^C isoleucine, ^13^C_2_ isoleucine, ^13^C_6_ isoleucine and ^13^C/^15^N isoleucine match expectation.

**Extended Data 5.**
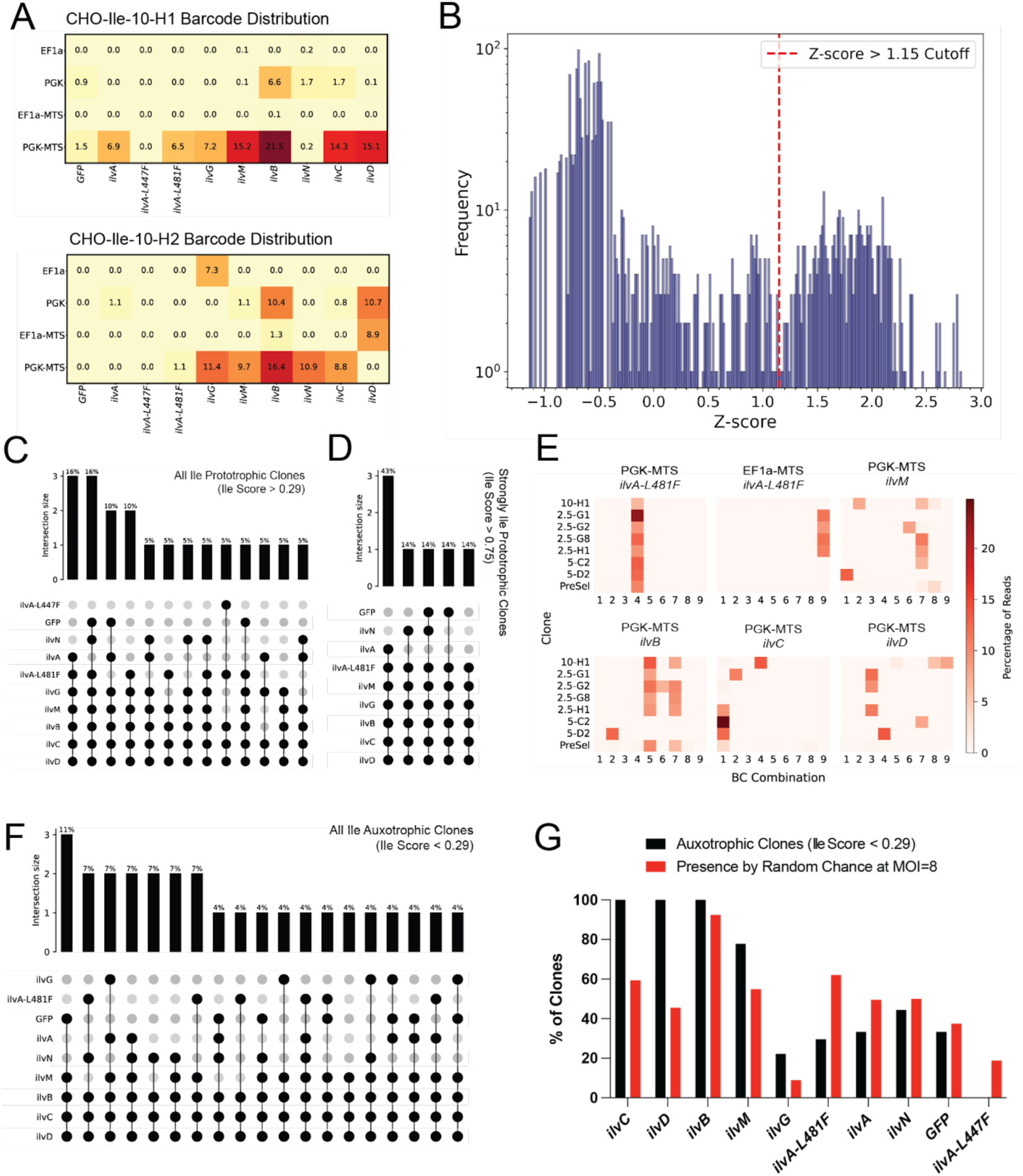
**Coding sequence usage underlying sequenced CHO clones selected on low isoleucine RPMI.** A. Normalized read counts for each TU within Clones CHO-Ile-10-H1 and CHO-Ile-10-H2. B. Log-transformed read counts were used to calculate a Z-score for each TU in each sample. A Z-score threshold of 1.15 was used to define TU presence vs. TU absence across the 46 clones. C. An UpsetPlot depicting which combinations of CDSes have been determined to be present within each of the 19 isoleucine prototrophic clones. D. An UpsetPlot depicting which combinations of CDSes have been determined to be present within each of the 7 strongly isoleucine prototrophic clones. E. Array of TUs determined to be involved in isoleucine prototrophic function and their 9 underlying barcode combinations (3 per promoter/OLS, 3 per CDS) across all strongly isoleucine prototrophic clones (Ile Score > 0.75). F. An UpsetPlot depicting which combinations of CDSes have been determined to be present within each of the 27 isoleucine auxotrophic clones. G. CDS usage amongst isoleucine auxotrophic clones is compared to the simulated expectation

**Extended Data 6.**
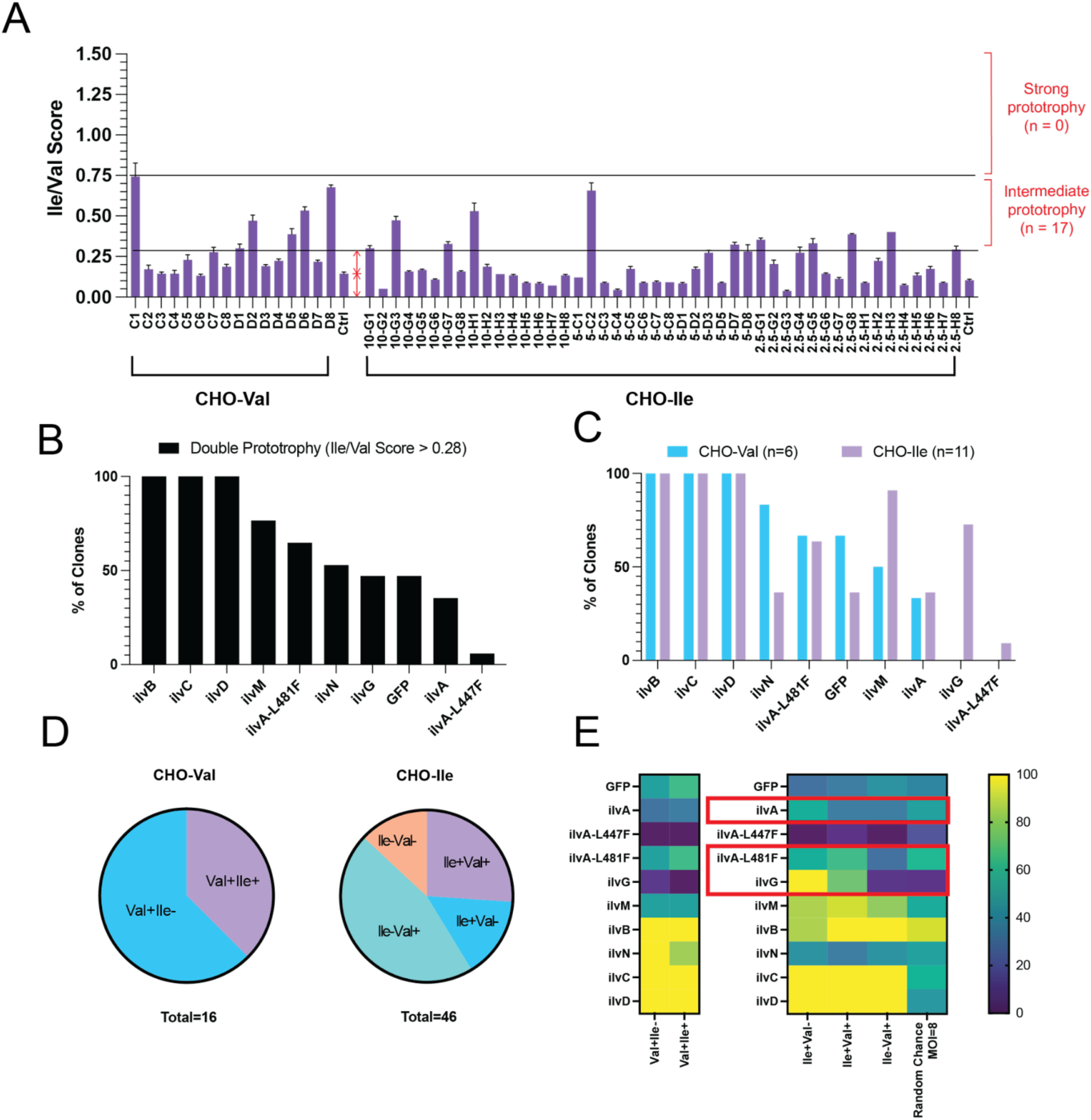
**Sampling CHO-Val and CHO-Ile clones for double isoleucine/valine prototrophy** A. Growth (PrestoBlue) measurements for 16 CHO-Val clones and 46 CHO-Ile clones cultured on isoleucine-free, valine-free RPMI. Triplicate wells were measured following 3 days on isoleucine-free, valine-free medium. Error bars represent standard deviation across triplicate wells. B. CDS presence amongst all double isoleucine/valine prototrophic clones. C. CDS presence amongst double isoleucine/valine prototrophic CHO-Val clones and double isoleucine/valine prototrophic CHO-Ile clones. D. Phenotype distribution amongst 16 CHO-Val clone and 46 CHO-Ile clones. E. CDS presence by phenotype in CHO-Val (left) and CHO-Ile clones (right) as well as the expected distribution according to simulated data. Notable differences across phenotypes among CHO-Ile clones are highlighted in red boxes for *ilvA, ilvA-L481F,* and *ilvG*.

**Extended Data 7.**
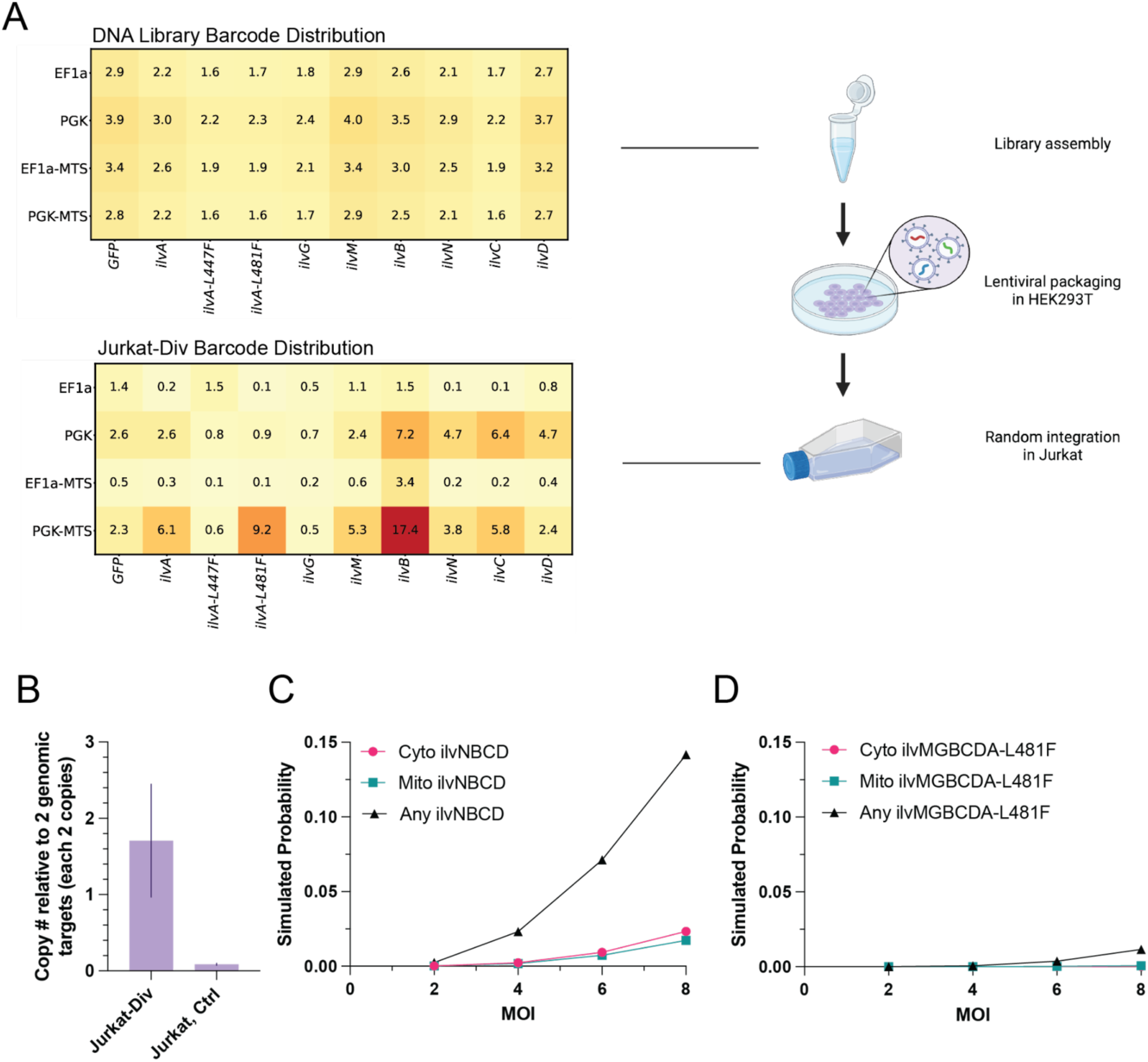
**Transcription unit distribution in Jurkat pre-infection (DNA Library) and post-infection (Jurkat-Div) of SGE library.** A. Transcription unit distribution prior to lentiviral packaging compared to transcription unit distribution after infection of Jurkat cells B. qPCR of TU integrants relative to two double copy genomic targets (RPP30, ALB) for estimation of MOI C. Probability of finding a presumed valine biosynthesis pathway combination post-infection based on a simulated infection of 3M Jurkat cells with the library distribution measured for Jurkat-Div cells in (A). Depicted is the probability of finding ilvN, ilvB, ilvC and ilvD irrespective of subcellular compartmentalization (black), all 4 genes localized to the cytoplasm (red), or all 4 genes localized to mitochondria (green) at a range of MOIs. D. Probability of finding a presumed isoleucine biosynthesis pathway combination post-infection based on a simulated infection of 3M Jurkat cells with the library distribution measured for Jurkat-Div cells in (A). Depicted is the probability of finding ilvM, ilvG, ilvB, ilvC, ilvD and ilvA-L481F irrespective of subcellular compartmentalization (black), all 6 genes localized to the cytoplasm (red), or all 6 genes localized to mitochondria (green) at a range of MOIs.

**Extended Data 8.**
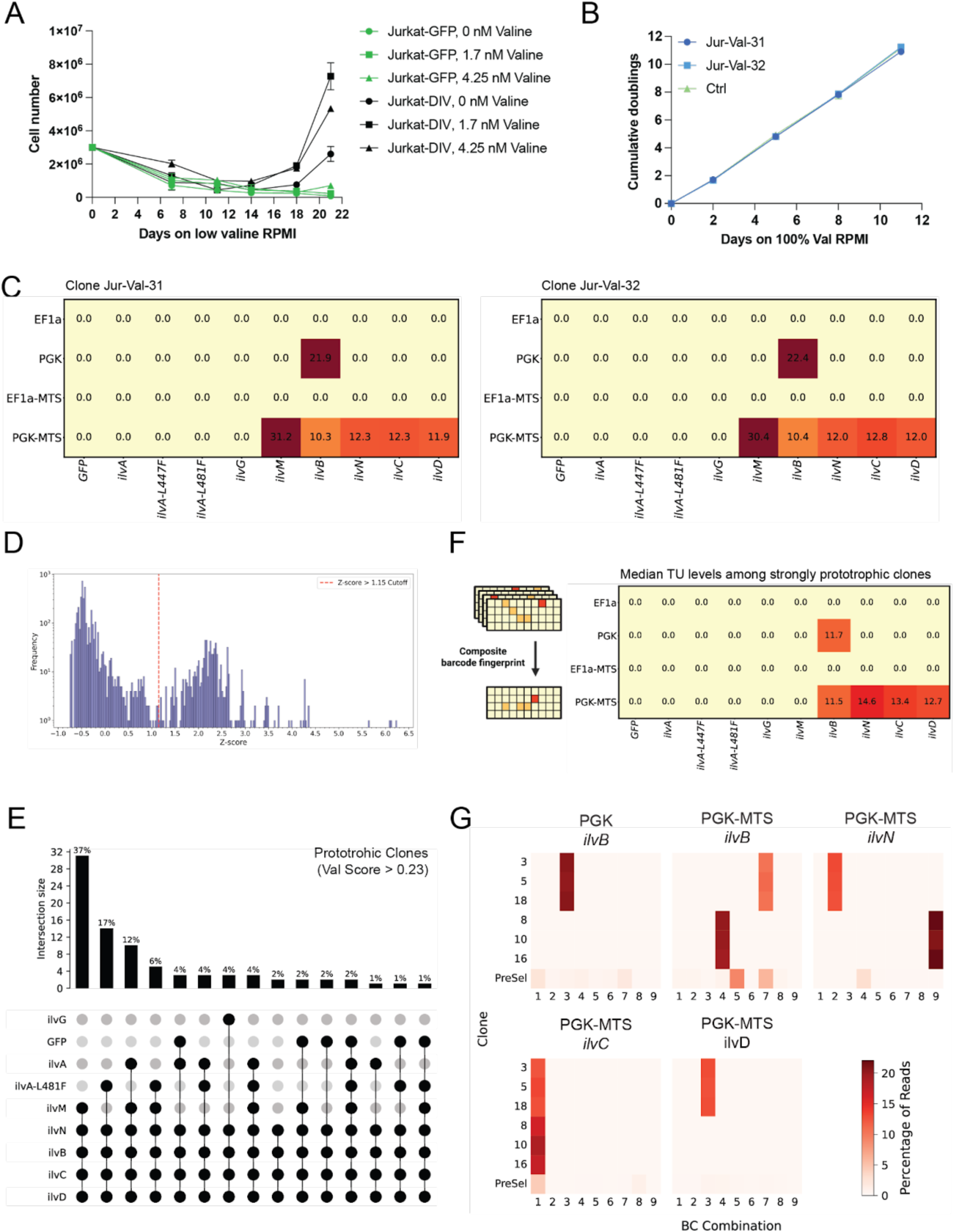
**Characterization of Jurkat-Div clones selected on low valine medium.** A. Cell counts of Jurkat-Div and Jurkat-GFP control cells over 21 days of selection on a range of low valine RPMI conditions. B. Growth curve of clones Jurkat-Val-31 and Jurkat-Val-32 cultured on valine-replete medium. Error bars represent standard deviation across triplicate wells. C. Normalized read counts for each TU, Clones Jurkat-Val-31 and Jurkat-Val-32 D. Log-transformed read counts were used to calculate a Z-score for each TU in each sample. A Z-score threshold of 1.15 was used to define TU presence vs. TU absence across the 119 clones. E. An UpsetPlot shows which combinations of CDSes have been determined to be present within each of the 84 valine prototrophic clones. F. Composite barcode ‘fingerprint’ comprised by median normalized read count across all strongly valine prototrophic clones (Ile Score > 0.75). G. Array of TUs determined to be involved in valine prototrophic function and their 9 underlying barcode combinations (3 per promoter/OLS, 3 per CDS) across all strongly isoleucine prototrophic clones (Val Score > 0.75). The array shows multiple groupings of matching barcode signatures indicating that several clones arose from the same founder clones.

**Extended Data 9.**
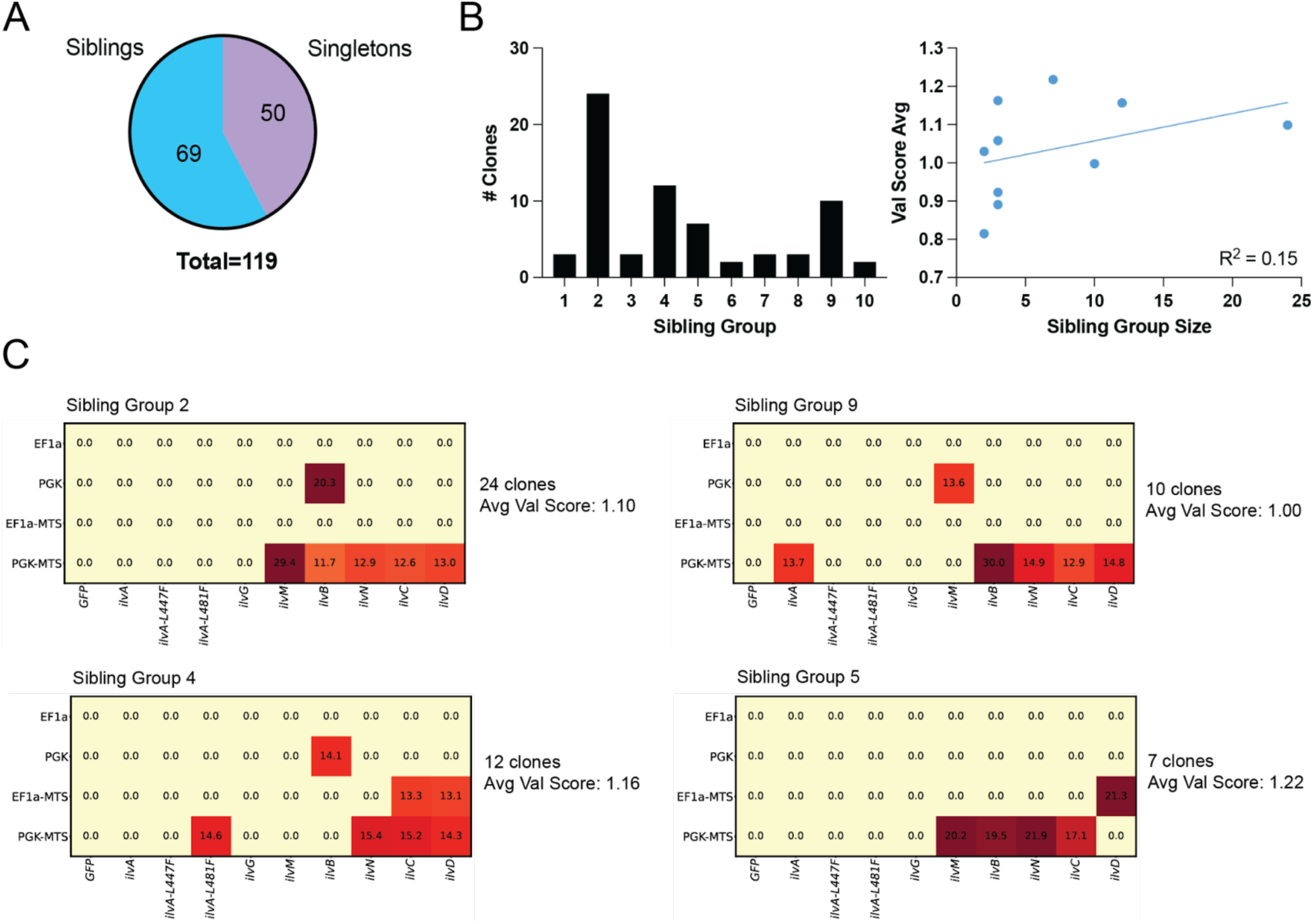
**Many clones arose from the same founder clones within the Jurkat-Val population** A. Distribution of clones that are part of sibling clonal groups and clones that are singletons. B. Graph-based clustering identified 10 separate sibling groups in Jurkat clones selected on low valine medium ranging in size from 2-24 clones each. Sibling group size does not correlate to an increase in average Val score. C. Representative heatmaps for each of the 4 largest sibling groups: Groups 2, 4, 9, and 5.

**Extended Data 10.**
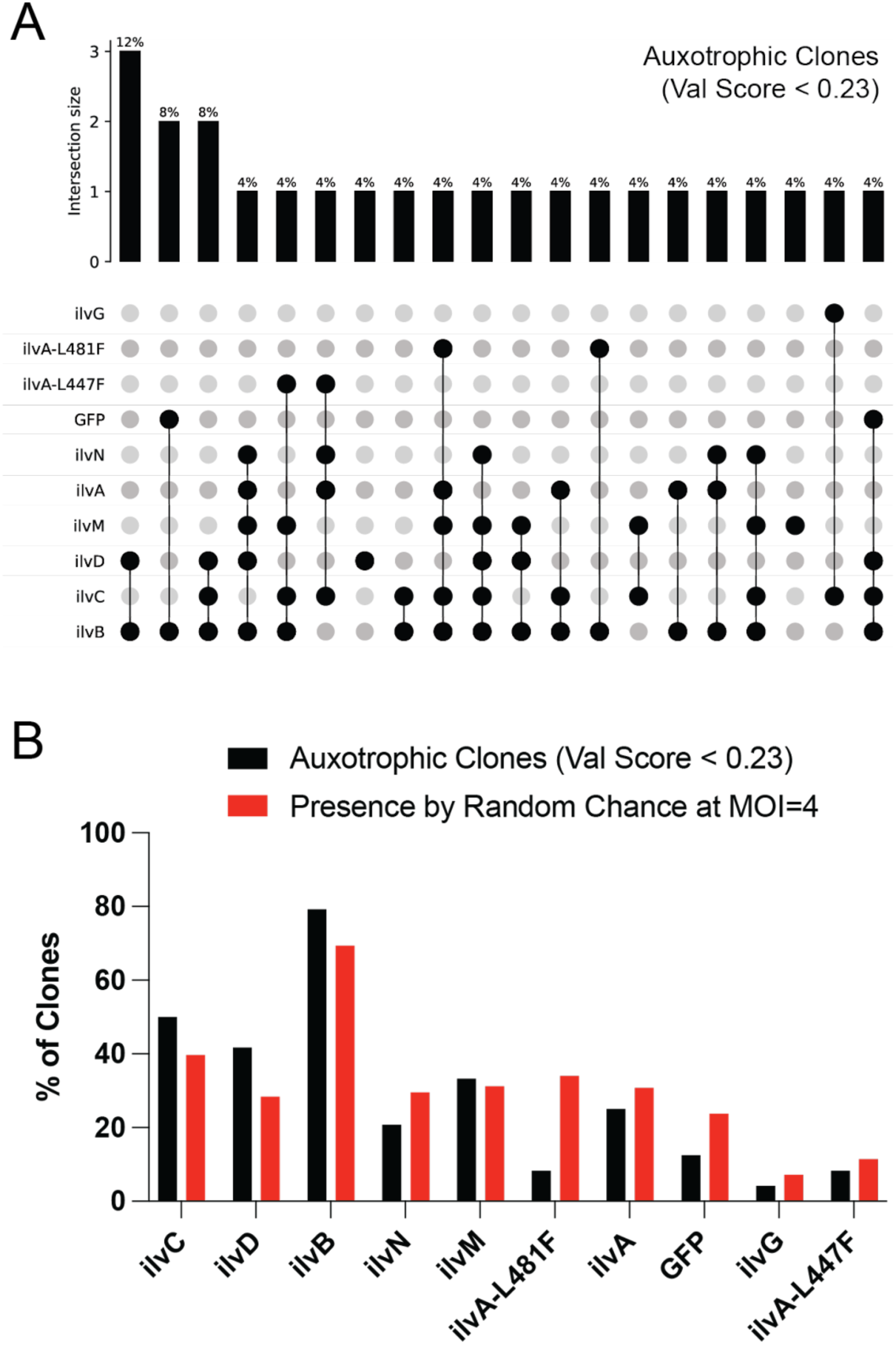
**Examining CDS usage in valine auxotrophic Jurkat clones selected on low valine medium.** A. An UpsetPlot depicting which combinations of CDSes have been determined to be present within each of 24 valine auxotrophic Jurkat clones carrying at least 1 TU integration. B. CDS usage amongst all valine auxotrophic clones carrying at least 1 TU integration is compared to the simulated expectation.

**Extended Data 11.**
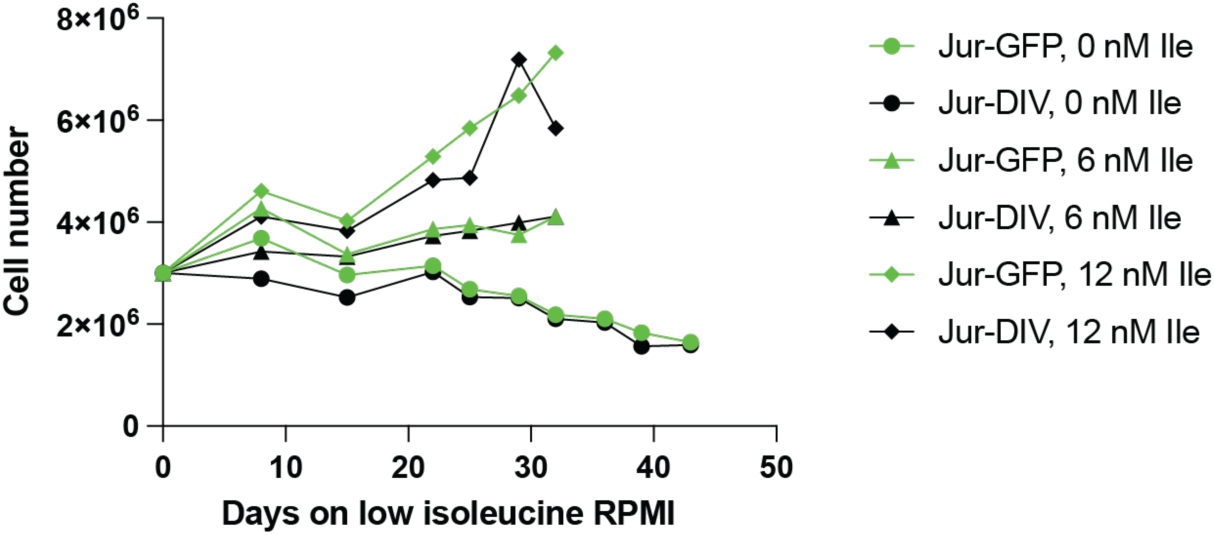
**Cell counts of Jur-Div cells subjected to low isoleucine medium selection over 32+ days alongside Jur-GFP control cells**

**Extended Data 12.**
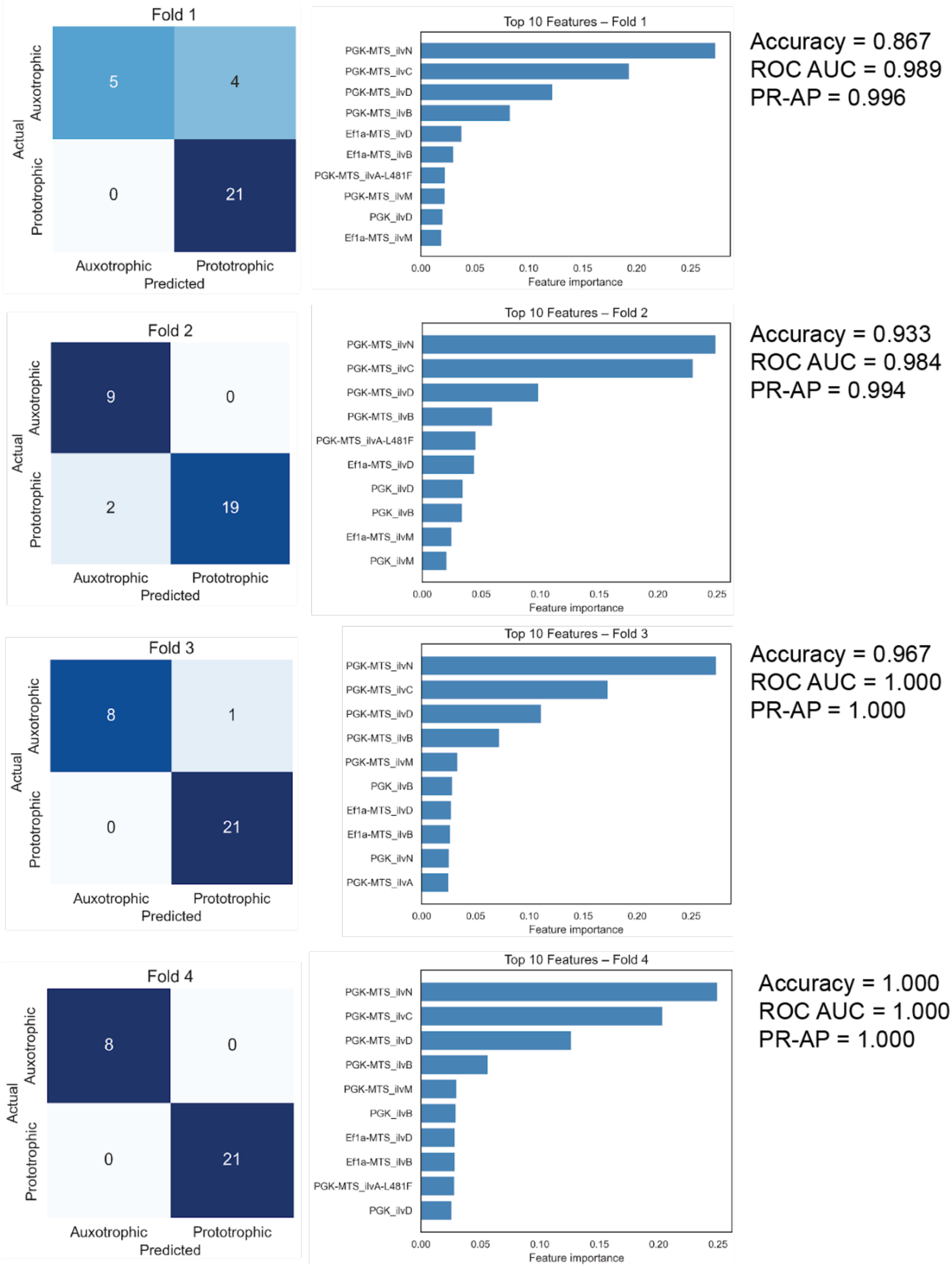
**4-fold cross-validation to validate Random Forest model stability across different train:test compositions. Classifications (left), top 10 features (middle) and metrics (right).**

